# Loss of a conserved disulfide bond defines Penetration2-related immune myrosinases in Brassicaceae

**DOI:** 10.64898/2026.08.10.742692

**Authors:** Gopal Singh, Himani Agrawal, Mariola Piślewska-Bednarek, Suthitar Singkaravanit-Ogawa, Chujia Jin, Anna Piasecka, Monidipa Bose, Sylwia Kuczewska, Aleksander Strugała, Łukasz Marczak, Milosz Ruszkowski, Yoshitaka Takano, Paweł Bednarek

## Abstract

- This study investigated whether PEN2/BGLU26 has uniquely evolved as an indole glucosinolate-hydrolysing myrosinase required for *Arabidopsis thaliana* pre-invasive immunity, or whether related myrosinases can replace its function when targeted to the same subcellular context.
- PEN2-homologous and other selected myrosinases from *A. thaliana* and *Brassica rapa* were expressed in the *pen2-*2 mutant background using a PEN2-like targeting strategy. The resulting lines were assessed by gene expression, protein accumulation, metabolite analysis and pathogen resistance assays. In parallel, targeted mutagenesis, structural comparison and phylogenetic analysis were used to examine molecular and evolutionary features of PEN2-related myrosinases.
- *At*BGLU27 and *Br*BABG.a, but not *At*BGLU18, *At*BGLU23 or *At*BGLU28, partially restored indole glucosinolate hydrolysis and resistance to *Colletotrichum tropicale* in *pen2-2*. Unlike *At*PEN2, both enzymes acted mainly constitutively and showed distinct substrate preferences. PEN2, BGLU27 and BABG proteins lacked conserved post-translational modification sites, including residues associated with a conserved disulfide bond. Restoring this disulfide bond in *At*PEN2 abolished its activity.
- PEN2-related myrosinases form an evolutionarily distinct BGLU lineage associated with indole glucosinolate metabolism in Brassicales. Loss of the conserved disulfide bond appears to be required for PEN2 activity, whereas additional PEN2-specific regulatory features are needed for pathogen-triggered, rather than constitutive, glucosinolate metabolism.

## Introduction

Plants produce a myriad of chemically diverse specialised metabolites in relation to their biotic and abiotic interactions. For instance, amino-acid-derived nitrogen and sulphur containing signature compounds in the members of Brassicaceae family, collectively referred as glucosinolates, are among the best studied specialised metabolites (Halkier & Gershenzon, 2006). Typically, glucosinolates are grouped into three broader categories based on their amino acid precursors: Met, Ala, Val, Leu or Ile-derived aliphatic glucosinolates; Phe-derived benzyl glucosinolates; and Trp-derived indolic glucosinolates (IGs) (Blažević *et al*., 2020). Constitutively, these thio-glucosides are biologically inactive and require specific β-thioglucosidases (also known as myrosinases or TGGs; ThioGlucoside Glucohydrolases) to release the unstable aglycones, which subsequently decompose into bioactive isothiocyanates, nitriles or other products (Blažević et al., 2020). Myrosinases represent Glycosyl Hydrolase Family 1 (GH1) β-glucosidases (BGLUs) and specifically hydrolyse thio-glucosidic bond to cleave glucose from glucosinolates (Sugiyama & Hirai, 2019). Early studies revealed that myrosinases are encoded in Brassicaceae plants by multigene families (Rask *et al*., 2000; Xu *et al*., 2004). For instance, six *TGG* genes have been identified in the model plant *A. thaliana* (*TGG1-6/BGLU34-39*), out of which four have been shown to encode functional myrosinases in the accession Col-0 (Andersson *et al*., 2009). While BGLUs possess typically two glutamic acids (EE) as catalytic residues in the initially studied myrosinases one of these residues, functioning as acid/base catalyst, was replaced with Gln (QE) suggesting this is a conserved feature distinguishing myrosinases from BGLUs hydrolysing *O*-glucosides (Burmeister *et al*., 1997; Rask *et al*., 2000; Xu *et al*., 2004). This conviction has been challenged with characterization of PENETRATION2 (PEN2)/BGLU26, which has two glutamic acids as catalytic residues and was identified as a myrosinase integrating IG metabolism into plant immune responses (Bednarek *et al*., 2009). It has been shown that *At*PEN2 constitutes an indispensable component of a metabolic immune pathway that controls entry of numerous filamentous pathogens, including *Blumeria graminis*, *Erysiphe pisi* and *Colletotrichum tropicale*, into *A. thaliana* epidermal cells (Lipka *et al*., 2005; Bednarek *et al*., 2009; Hiruma *et al*., 2010). This function has been shown to be dependent on subcellular localization of this protein. PEN2 is anchored in the mitochondrial membranes and re-localizes, together with mitochondria, to the sites of attempted pathogen entry where it likely generates high local concentrations of IG-hydrolytic products (Lipka *et al*., 2005; Fuchs *et al*., 2015). This unique localization is dependent on the 65 aa long unique C-terminal tail containing a 21 aa long predicted transmembrane domain, which has been shown to act as tail anchor (TA) inserting in the outer mitochondrial membrane (Fuchs *et al*., 2015).

Among PEN2 products, indol-3-ylmethyl amine (I3A) and raphanusamic acid (RA) have been shown to be formed from indol-3-ylmethyl glucosinolate (I3G) in response to pathogen inoculation or flagellin-epitope (flg22) treatment (Fig. 1) (Bednarek *et al*., 2009; Frerigmann *et al*., 2016). In addition to triggering PEN2-mediated I3G metabolism, pathogen recognition specifically induces expression of *CYP81F2* encoding P450 monooxygenase, which hydroxylates I3G to 4-hydroxy-I3G (4OHI3G) (Clay *et al*., 2009). This derivative can be subsequently methylated by IG-*O*-methyltransferases (IGMT1-4) to produce 4-methoxy-I3G (4MI3G; Fig. 1) (Pfalz *et al*., 2011). It has been shown that *pen2* mutants hyper-accumulate 4MI3G upon pathogen challenge or flg22 treatment, suggesting PEN2 also hydrolyses 4OH/4MI3G in addition to I3G (Bednarek *et al*., 2009; Clay *et al*., 2009; Frerigmann *et al*., 2016). Moreover, pathogen assays with single and double *pen2* and *cyp81f2* mutants revealed that both enzymes act on the same immune pathway to restrict entry of pathogens (Bednarek et al., 2009). This indicated that although PEN2 can metabolize I3G and 4OH/4MI3G, only product(s) of hydrolysis of the substituted IGs are important to restrict pathogen entry. 4-*O*-β-D-glucosyl-indol-3-yl formamide (4OGlcI3F) has been identified as one of the products of PEN2-mediated 4OH/4MI3G hydrolysis, however, function of this compound in penetration resistance remains obscure (Fig. 1) (Lu *et al*., 2015).

**Figure 1.**
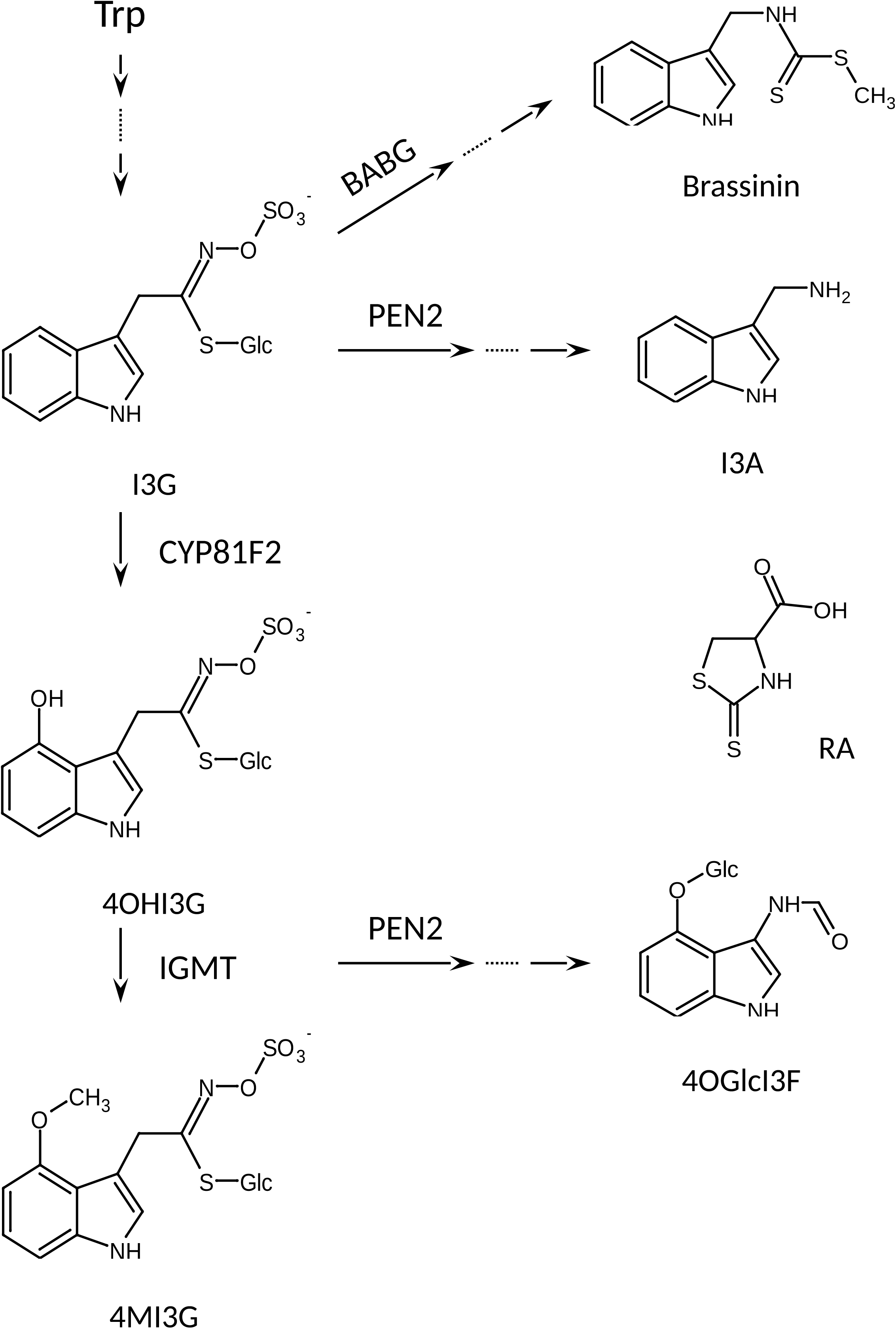
Contribution of Penetration2 (PEN2/BGLU27) and Brassinin associated β-glucosidase (BABG) to pathogen triggered indole glucosinolate metabolism. I3G - indol-3-ylmethylglucosinolate; 4MI3G - 4-methoxy-I3G; I3A - indol-3-ylmethylamine; RA - raphanusamic acid; 4OGlcI3F - 4-*O*-Glc-indol-3-yl formamide. Dashed lines indicate multiple enzymatic steps.

As mentioned above, discovery of PEN2 myrosinase activity challenged the belief that these enzymes have unique QE catalytic residues. A later study revealed that one of the *At*PEN2 close homologs, *At*BGLU23/PYK10, also has myrosinase activity despite its EE signature (Nakano *et al*., 2017; Yamada *et al*., 2020). This enzyme is the main proteinaceous component of root endoplasmic reticulum (ER) bodies and is postulated to provide resistance against biotic stress (Wilkens *et al*., 2026). Detailed sequence analysis of *At*BGLUs revealed that *At*BGLU18-33, forming a clade neighbouring the TGG clade in the BGLU phylogenetic tree, possess unique conserved positively charged arginine/lysine (K/R) residues (Fig. S1). According with structure modelling of these BLUSs and TGGs, as well as with the earlier structural study of a TGG from *Sinapsis alba* these K/R residues could contribute to the binding of the negatively charged glucosinolate molecule in the substrate binding pocket (Burmeister *et al*., 1997; Nakano *et al*., 2017). Based on these findings, *At*BGLU18-33 have been postulated to possess myrosinase activity and proposed to be referred as EE-myrosinases, to distinguish them from the investigated earlier QE-myrosinases (TGGs) (Nakano *et al*., 2017). This hypothesis has been partially validated with studies reporting function of *At*BGLU28 and *At*BGLU30 in sulfur homeostasis by glucosinolate hydrolysis in *A. thaliana* (Zhang *et al*., 2020; Sugiyama *et al*., 2021). Some indirect evidence pointed also at possible myrosinase activity of *At*BGLU18 and *At*BGLU21, which similar to *At*BGLU23, localize to ER-bodies in *A. thaliana* (Nakazaki *et al*., 2019; Yamada *et al*., 2020). Apart from these enzymes characterized in *A. thaliana*, *Brassica* species have been shown to possess specific EE-myrosinases, homologous with *At*BGLU18-33, called brassinin-associated β-glucosidases (BABGs) capable to hydrolyse IGs for the biosynthesis of sulphur-containing phytoalexin brassinin and its derivatives (Fig. 1) (Klein & Sattely, 2017).

In the present study, we propose the question whether PEN2 has specifically evolved as the unique myrosinase integrating IGs metabolism in immune responses or other EE type myrosinasess can complement its function in *pen2* mutant background if produced under identical subcellular localization. Our findings shed light on several key evolutionary attributes of plant immunity related myrosinases in Brassicaceae family.

## Material and methods

### Plant material

We used *A. thaliana* Col-0, *pen2-1*, and *pen2-2* (GABI-KAT 134C04) lines described previously (Lipka *et al*., 2005). Seeds of the investigated lines were sown in soil and germinated under short-day conditions (22 °C; 10 h light, 14 h darkness). Seedlings of transgenic plants were selected with glufosinate-ammonium spray (100 mg/L; Cayman Chemical, Ann Arbor, MI). After 2 weeks, seedlings intended for gene expression, western blot, and metabolic analyses were transferred to Jiffy-7 pellets (Jiffy Group, Zwijndrecht, Netherlands) and grown for additional 3 weeks under the same conditions. To induce the PEN2 pathway, plants were sprayed with 5 µM synthetic peptide corresponding to the 22-amino acid epitope of bacterial flagellin (flg22; ProteoGenix, Schiltigheim, France) in 0.1% (v/v) Tween 20 (Sigma-Aldrich, St. Louis, MO) (Felix *et al*., 1999). Control plants were sprayed with sterile water containing 0.1% (v/v) Tween 20. Leaf samples were collected 24 h after treatment, immediately frozen in liquid nitrogen, and stored at −80 °C until use. To obtain plants for transformation, 2-week-old seedlings were transferred to soil-filled pots and grown for an additional 4 weeks under short-day conditions. Plants were then moved to long-day conditions (22 °C; 16 h light, 8 h darkness) to induce flowering.

### Construct preparation and generation of transgenic lines

Plasmids for plant transformation were generated based on the binary vector backbone of pAMPAT-MCS (AY436765). We used a pAMPAT vector variant containing the precloned *AtPEN2* (*AT2G44490*) cDNA coding sequence flanked by 1213 bp of the *AtPEN2* 5′ untranslated region and the 35S terminator (Fig. S3A) (Lipka *et al*., 2005). This construct was also used to obtain control lines expressing *AtPEN2* in *pen2-2* mutant background. To enable cloning of all other gene/cDNA fragments, we used the *Hin*dIII restriction site located between the promoter and the *AtPEN*2 cDNA sequence, together with the *Aat*II restriction site located in the 5′ end of the *AtPEN2-TA* CDS, to remove the core *At*PEN2 coding sequence (Fig. S3A). This fragment was replaced with a synthetic DNA insert containing *Asi*SI and *Sbf*I restriction sites, followed by 1 bp preventing the proper reading frame and 14 bp of the 5′ *At*PEN2-TA CDS located upstream of the *Aat*II site. The synthetic insert was generated by annealing two phosphorylated oligonucleotides designed to produce *Hin*dIII- and *Aat*II-compatible sticky ends, respectively (Table S1). Using the introduced *Asi*SI and *Sbf*I sites in this modified plasmid, we cloned all sequences of interest downstream of the *AtPEN2* promoter and in frame with the *AtPEN2-TA* CDS. For cloning *AtBGLU18* (*AT1G52400*) and *AtBGLU27* (*AT3G60120*), the truncated regions of respective genes were amplified from genomic DNA of Col-0 using gene specific primers (Fig. S3B; Table S1). Sequence encoding 15 aa N-terminal of *At*PEN2 was added in *AtBGLU18* through forward primer. For generating *BrBABG.a* (*Brara.D02695*), *AtBGLU23* (*AT3G09260*) and *AtBGLU28* (*AT2G44460*) chimeric genes, respective CDS regions were synthesized as synthetic genes (Synbio Technologies, Monmouth Junction, NJ) and cloned into the recipient vector using *Asi*SI and *Sbf*I restriction sites. Mutagenesis of *AtPEN2* cDNA to replace V^211^ encoding codon with codons for CY was performed with NEB Q5® Site-Directed Mutagenesis Kit (New England Biolabs, Ipswich, MA) (Table S1).

Sanger sequencing-confirmed constructs were introduced into *Agrobacterium tumefaciens* GV3101 (pMP90RK) by electroporation (Zhang *et al*., 2006). The respective *A. tumefaciens* strains were then used for floral dip transformation of *pen2-2* plants. The resulting T_1_ plants were selected with glufosinate-ammonium spray (100 mg/L; Cayman Chemical). Individual selected plants were screened for expression of the respective transgenes. Plants showing relatively high transgene expression were kept for seed production and further characterization in the subsequent generations.

### RNA extraction and qPCR analysis for gene expression

For monitoring gene expression total RNA was extracted from the leaf samples using RNeasy Plant Mini Kit (Qiagen, Hilden, Germany), followed by cDNA synthesis using Omniscript RT Kit (Qiagen). Quantitative real-time PCR (qRT-PCR) analysis was performed with iTag Universal SYBR Green Supermix (Bio-Rad, Hercules, CA) using the CFX Connect Real-Time System (Bio-Rad). *Actin1* (*At2G37620*) was selected as the reference gene, while primer pair specific for the sequence encoding *At*PEN-TA was used for monitoring relative expression of *AtPEN2* and all investigated transgenes (Table S1). Quantification of relative gene expression was performed using 2^−ΔΔCt^ method (Livak & Schmittgen, 2001).

### Protein extraction and western blot analysis

Plant tissue was grinded with mortar and pestle in liquid nitrogen. Protein extraction was performed by adding lysis buffer (20 mM HEPES; pH 7.5, 13% sucrose, 1mM EDTA, 1M DTT) with cOmplete^™^ Mini Protease Inhibitor Cocktail (Merck Life Sciences, Darmstadt, Germany) to the grinded tissue. Tissue homogenates were centrifuged, supernatant was collected and the protein concentration was quantified with standard Bradford assay. Proteins were separated with SDS-PAGE electrophoresis on 8% separating gel, transferred to a poly-vinylidene fluoride membrane (Merck Life Sciences). The *At*PEN2-TA antiserum was generated against the peptide ^506^EEKKESYGKQLLHSVQ^521^ in rabbit, affinity-purified (Eurogentec, Seraing, Belgium) (Lipka *et al*., 2005). We used horseradish-conjugated anti-rabbit IgG (Cell Signaling Technology, Danvers, MA) as secondary antibody. General blotting and detection procedures were as described earlier (Bednarek *et al*., 2009).

### Metabolite extraction and analysis

Frozen leaf samples (∼200 mg) were homogenized in DMSO (2.5 μl per 1 mg fresh weight), centrifuged, and the supernatants were collected for further analysis. Detection and quantification of IGs, I3A, and RA were performed by high-performance liquid chromatography (HPLC) with UV and fluorescence detection. Extracts were analysed using an UltiMate 3000 RS HPLC system equipped with diode-array (DAD) and fluorescence (FLD) detectors (Thermo Fisher Scientific, Waltham, MA). Samples were separated on two Atlantis T3 C18 columns connected in series (150 × 2.1 mm and 50 × 2.1 mm, 3 µm; Waters, Milford, MA) using 0.1% trifluoroacetic acid (TFA) as solvent A and 98% acetonitrile (ACN)/0.1% TFA as solvent B, at a flow rate of 0.25 ml/min and 20 °C. The following solvent A gradient was used: 100% at 0 min, 100% at 2 min, 90% at 9 min, 72% at 30 min, 50% at 33 min, 20% at 40 min, and 100% at 41 min.

I3G, 4MI3G, RA and 6-*O*-Glc-indole-3-carboxylic acid (6OGlcICA) were monitored and quantified based on UV chromatograms recorded at 273 nm. I3A was quantified using FLD chromatograms recorded at 275/350 nm (excitation/emission). Peaks of metabolites of interest were identified and quantified based on analysis of the respective standards: I3G, 4MI3G and 6OGlcICA purified from plant tissue, synthetic I3A, and RA from Sigma-Aldrich (Bednarek *et al*., 2005; Bednarek *et al*., 2009).

Due to the relatively low concentration of 4OGlcI3F in the analysed extracts, liquid chromatography coupled with mass spectrometry (LC-MS) was used for reliable identification and quantification of this compound. LC-MS analysis was performed using an UltiMate 3000 RS system (Thermo Fisher Scientific) coupled to a TIMS-TOF mass spectrometer (Bruker Daltonics, Hamburg, Germany) in positive ionization mode, under conditions described previously (Piślewska-Bednarek *et al*., 2026). The 4OGlcI3F signal was identified based on analysis of the available synthetic standard (Lu *et al*., 2015).

### Fungal strain and culture conditions

The fungal pathogen *Colletotrichum tropicale* (*Ctro*; previously designated as *Colletotrichum gloeosporioides* S9275) was kindly provided by Dr. Shigenobu Yoshida from the National Institute for Agro-Environmental Sciences, Japan. The strain was maintained on 2.5% (w/v) potato dextrose agar (PDA; Difco, Detroit, MI, USA). Culturing was performed at 24°C under a 16-h photoperiod using black light (FS20S/BLB 20W; Toshiba, Tokyo, Japan) followed by an 8-h dark period.

### Pathogen inoculation, lesion development analysis and fungal invasion assay in *A. thaliana*

Seeds of investigated lines were sown on Grodan rockwool (Rockwool B.V., Roermond, Nederland), kept at 4°C in the dark for 2 days for stratification, and then grown at 22–23°C under a 16-h light/8-h dark photoperiod. Transgenic *At*BGLU27 and *Br*BABG plants were selected by spraying seedlings three times with a 1/2000 dilution of BASTA (glufosinate-ammonium; BASF, Japan). Confirmed positive transformants were subsequently used for *Ctro* inoculation.

To evaluate lesion development, each 4 weeks plant leaves were drop-inoculated with a 5 µL droplet of *Ctro* conidial suspension (2.5 × 10^5^ conidia/mL) supplemented with 0.1% (w/v) glucose. Following inoculation, plants were incubated at 22-23°C under a 16-h light/8-h dark at 100% relative humidity. The photographs were taken at 5 dpi. Disease lesions were imaged and measured quantitatively using ImageJ software (http://imagej.net). For the fungal invasion assay, 14 days cotyledons were inoculated by applying a 2 µL droplet of the same conidial suspension (2.5 × 10^5^ conidia/mL). The invasive hyphae were observed and analyzed via light microscopy at 14 hours post-inoculation (hpi).

### Design of *At*PEN2 mutant with reintroduced disulfide bond (PEN2_DSB)

The mutant was designed by comparing the *At*PEN2 structural model retrieved from AlphaFoldDB (Fleming *et al*., 2025) with the experimentally determined structure of *S. alba* myrosinase (PDB ID: 1MYR) (Burmeister *et al*., 1997). This comparative approach was used because AlphaFold2 (the version available at the time of design) does not explicitly model DSBs.

### Proteomic analysis of DSB formation

Protein extracts were prepared in the non-reducing variant of the lysis buffer described above, without DTT. After SDS-PAGE, gel fragments corresponding to the expected molecular weight of *At*PEN2 variants, approximately 60 kDa, were excised with a scalpel, cut into approximately 1 × 1 mm squares, and transferred into low-binding microcentrifuge tubes. Gel pieces were washed with 10 mM ammonium bicarbonate/50% ACN to remove buffer components, dehydrated with ACN, rehydrated with 10 mM ammonium bicarbonate, washed again with ACN, and dried in a vacuum concentrator. Proteins were digested under non-reducing conditions to preserve DSBs and avoid cysteine alkylation. Dried gel pieces were rehydrated with sequencing-grade trypsin in 10 mM ammonium bicarbonate, using the minimal volume required to cover the gel pieces, and incubated for 60 min at 4 °C. A small volume of 10 mM ammonium bicarbonate was then added to keep the gel pieces wet, and digestion was carried out overnight at 37 °C. Peptides were extracted with 50% ACN containing 1% TFA, assisted by 10 min sonication. Combined extracts were dried in a vacuum concentrator and reconstituted in 0.1% formic acid (FA) before analysis.

Proteomic analysis was performed using a Dionex UltiMate 3000 RSLC nanoLC system coupled to an Orbitrap Exploris 480 mass spectrometer (Thermo Fisher Scientific). Peptides were separated on an Acclaim PepMap RSLC nanoViper C18 reversed-phase analytical column (75 µm × 25 cm, 2 µm particle size; Thermo Fisher Scientific) maintained at 30 °C. Chromatographic separation was performed at 300 nL/min using mobile phases containing 0.1% FA, with peptide elution over a 190 min linear ACN gradient from 4% to 60%.

### Identification of BGLU homologs and phylogenetic analyses

Protein sequences of close *At*PEN2, *At*BGLU27, and *Br*BABG.a homologs from the majority of selected species (Table S2) were retrieved from Phytozome and other genomic databases using local BLAST searches (Goodstein *et al*., 2012; Kagale *et al*., 2014; Willing *et al*., 2015; Gan *et al*., 2016; Nguyen *et al*., 2019). In case of *Tarenaya hassleriana*, *Gynandropsis gynandra* and *Capparis spinosa* the whole proteome files were downloaded from respective repositories and searched for homologous sequences (Cheng *et al*., 2013; Wang *et al*., 2022; Hoang *et al*., 2023). The selected protein sequences were aligned using the MUSCLE algorithm, and a maximum-likelihood phylogenetic tree was generated with MEGA 12 software in adaptive mode with threshold 1 (Kumar *et al*., 2024).

## Results

### *At*BGLU27 and *Br*BABG.a complement myrosinase activity in *pen2-2*

To check if any other EE type myrosinase can replace *At*PEN2 function in pre-invasive resistance we expressed genes encoding selected representatives of these enzymes in *pen2-2* mutant background driven by the native *PEN2* promoter. Selected enzymes included *At*BGLU27 and *Br*BABG.a, which are closest homologs of *At*PEN2 (Fig. S1). Among these, *Br*BABG.a has been unambiguously shown to possess myrosinase activity and contribute to brassinin biosynthesis, while *AtBGLU27* expression has been shown to be upregulated with immune responses, however, its function remains obscure (Fig. 1)(Iven *et al*., 2012; Klein & Sattely, 2017; Souza *et al*., 2017). To express these selected BGLUs in PEN2 alike cellular and sub-cellular condition, we considered to generate chimeric *BGLU* genes. Firstly, we truncated sequences encoding the N- and C-terminal regions that could act as sorting/retention signals or TAs, and secondly, we added a sequence encoding the 65 aa C-terminal TA of *At*PEN2 (PEN2-TA) fragment for anchoring of chimeric proteins into mitochondrial membrane, which is essential for PEN2 immune function (Table 1; Fig. S2, S3)(Fuchs *et al*., 2015).

**Table 1.** Key attributes, including predicted post-translational modifications, of *Arabidopsis thaliana* BGLUs based on UniProt database (UniProt Consortium, 2025). Presence of signal peptide according to Xu *et al*. 2004. *Br*BABG protein sequences were retrieved from *Brassica rapa* genome from Phytozome and analysed for conservation of respective amino acid residues with MEGA 12 software (Goodstein *et al*., 2012; Kumar *et al*., 2024).

| BGLU | Sequence ID | Uniprot ID | Signal peptide | Disulfide bonds | N-Glycosylation sites | ER retention signal |
| --- | --- | --- | --- | --- | --- | --- |
| BGLU1 | At1g45191 | Q3ECW8 | + | 210-217 | 216, 221, 441, 473, 512 | - |
| BGLU2 | At5g16580 | Q9FMD8 | + | 69-72 | 71, 76, 222, 290 | - |
| BGLU3 | At4g22100 | O65458 | + | 203-210 | 64, 209, 214, 361, 429, 461, 485, 500 | - |
| BGLU4 | At1g60090 | Q9ZUI3 | + | 205-212 | 211, 216, 436, 468 | - |
| BGLU5 | At1g60260 | Q8RXN9 | + | 205-212 | 216, 361, 424, 456, 495 | - |
| BGLU6 | At1g60270 | Q682B4 | + | 206-213 | 217, 362 | - |
| BGLU7 | At3g62740 | Q9LZJ1 | + | - | 208, 359, 425, 457, 479 | - |
| BGLU8 | At3g62750 | Q67XN2 | + | - | 65, 202, 354, 452, 474, 490 | - |
| BGLU9 | At4g27820 | Q9STP4 | + | 204-212 | 211, 216, 363, 429, 461, 483, 499 | - |
| BGLU10 | At4g27830 | Q93ZI4 | + | 207-215 | 214, 219, 365, 431, 463, 485, 501 | - |
| BGLU11 | At1g02850 | B3H5Q1 | + | 209-217 | 216, 221, 364, 388 | - |
| BGLU12 | At5g42260 | Q9FH03 | + | 219-227 | 81, 226, 358 | - |
| BGLU13 | At5g44640 | Q9LU02 | + | 219-227 | 81, 226, 358 | - |
| BGLU14 | At2g25630 | Q9SLA0 | + | 218-226 | 80, 225, 357 | - |
| BGLU15 | At2g44450 | O64879 | + | 219-227 | 24, 25, 81, 226 | - |
| BGLU16 | At3g60130 | Q9M1D0 | + | 218-226 | 80, 357 | - |
| BGLU17 | At2g44480 | O64882 | + | 223-230 | 229, 361, 371, 510 | - |
| BGLU18 | At1g52400 | Q9SE50 | + | 226-237 | 189, 466, 499 | 525-528 (REEL) |
| BGLU19 | At3g21370 | Q9LIF9 | + | 220-231 | 183, 462, 495 | 524-527 (HEEL) |
| BGLU20 | At1g75940 | Q84WV2 | + | 224-235 | 187, 468, 501 | 532-535 (HDEL) |
| BGLU21 | At1g66270 | Q9C525 | + | 223-230 | 61, 494 | 521-524 (RDEL) |
| BGLU22 | At1g66280 | Q9C8Y9 | + | 223-230 | 61, 494 | 521-524 (KDEL) |
| BGLU23 | At3g09260 | Q9SR37 | + | 222-230 | 60, 461, 494 | 521-524 (KDEL) |
| BGLU24 | At5g28510 | Q9LKR7 | + | 226-239 | 64, 88, 388, 437, 442, 470, 503 | 530-533 (KDEL) |
| BGLU25 | At3g03640 | O82772 | + | 222-230 | - | - |
| BGLU26 | At2g44490 | O64883 | - | - | - | - |
| BGLU27 | At3g60120 | Q9M1D1 | - | - | - | - |
| BrBABG.a | Brara.D02695 | - | - | - | - | - |
| BrBABG.b | Brara.E00436 | - | - | - | - | - |
| BGLU28 | At2g44460 | Q4V3B3 | + | 216-224 | 255, 330, 370, 430, 521, 544 | - |
| BGLU29 | At2g44470 | Q8GXT2 | + | 216-224 | 255, 331, 371, 522, 553 | - |
| BGLU30 | At3g60140 | Q9M1C9 | + | 213-221 | 328, 368, 524, 544 | - |
| BGLU31 | At5g24540 | Q9FLU9 | + | 219-227 | 68, 374, 425 | - |
| BGLU32 | At5g24550 | Q9FLU8 | + | 219-227 | 68, 374, 425 | - |
| BGLU33 | At2g32860 | O48779 | + | 282-290 | 344, 419, 432, 439, 491 | - |
| BGLU34 | At1g47600 | Q8GRX1 | + | 31-450, 39-445, 230-233 | 46, 53, 428, 489 | - |
| BGLU35 | At1g51470 | Q3ECS3 | + | 31-450, 39-445, 230-233 | 46, 53, 222, 428, 489 | - |
| BGLU36 | At1g51490 | Q9C8K1 | - | 204-207 | 28, 260, 462 | - |
| BGLU37 | At5g25980 | Q9C5C2 | + | 36-460, 44-456, 236-244 | 340, 384, 504 | - |
| BGLU38 | At5g26000 | P37702 | + | 24-449, 32-445, 224-232 | 33, 108, 175, 236, 356, 379, 493, 512 | - |
| BGLU39 | At5g48375 | Q3E8E5 | + | - | 33, 336 | - |
| BGLU40 | At1g26560 | Q9FZE0 | + | 217-225 | 229, 278, 349, 507 | - |
| BGLU41 | At5g54570 | Q9FIU7 | + | 216-224 | 118, 445, 489 | - |
| BGLU42 | At5g36890 | Q9FIW4 | - | - | 420 | - |
| BGLU43 | At3g18070 | Q9LV34 | + | 213-220 | 77, 219, 360, 416, 424, 448 | - |
| BGLU44 | At3g18080 | Q9LV33 | + | 224-231 | 86, 230, 427 | - |
| BGLU45 | At1g61810 | O80689 | + | 220-227, 352-357 | 3, 226, 378, 435 | - |
| BGLU46 | At1g61820 | O80690 | + | 217-224, 349-354 | 223, 375 | - |
| BGLU47 | At4g21760 | Q9SVS1 | + | 240-247, 371-376 | 93, 246, 419, 432 | - |

After obtaining stable transgenic lines, we analysed the expression of *AtBGLU27* and *BrBABG.a* chimeric genes in leaves of respective transgenic plants using primers specific for *AtPEN2-TA* (Table S1). For *AtBGLU27* (lines #1, 2, 3) the expression levels were found to be similar or even higher as compared to *AtPEN2* in Col-0. Despite screening a similar number of T_1_ plants, in case of *BrBABG.a* we only manage to isolate lines with relatively lower transgene expression (Fig. 2A). We then analysed the protein abundance of these chimeric BGLUs using an antibody specific to a peptide from *At*PEN2-TA. Complementing gene expression pattern, the signal abundance recorded for *At*BGLU27 was similar to *At*PEN2 in Col-0, while comparatively low in case of *Br*BABG.a (Fig. 2B).

**Figure 2.**
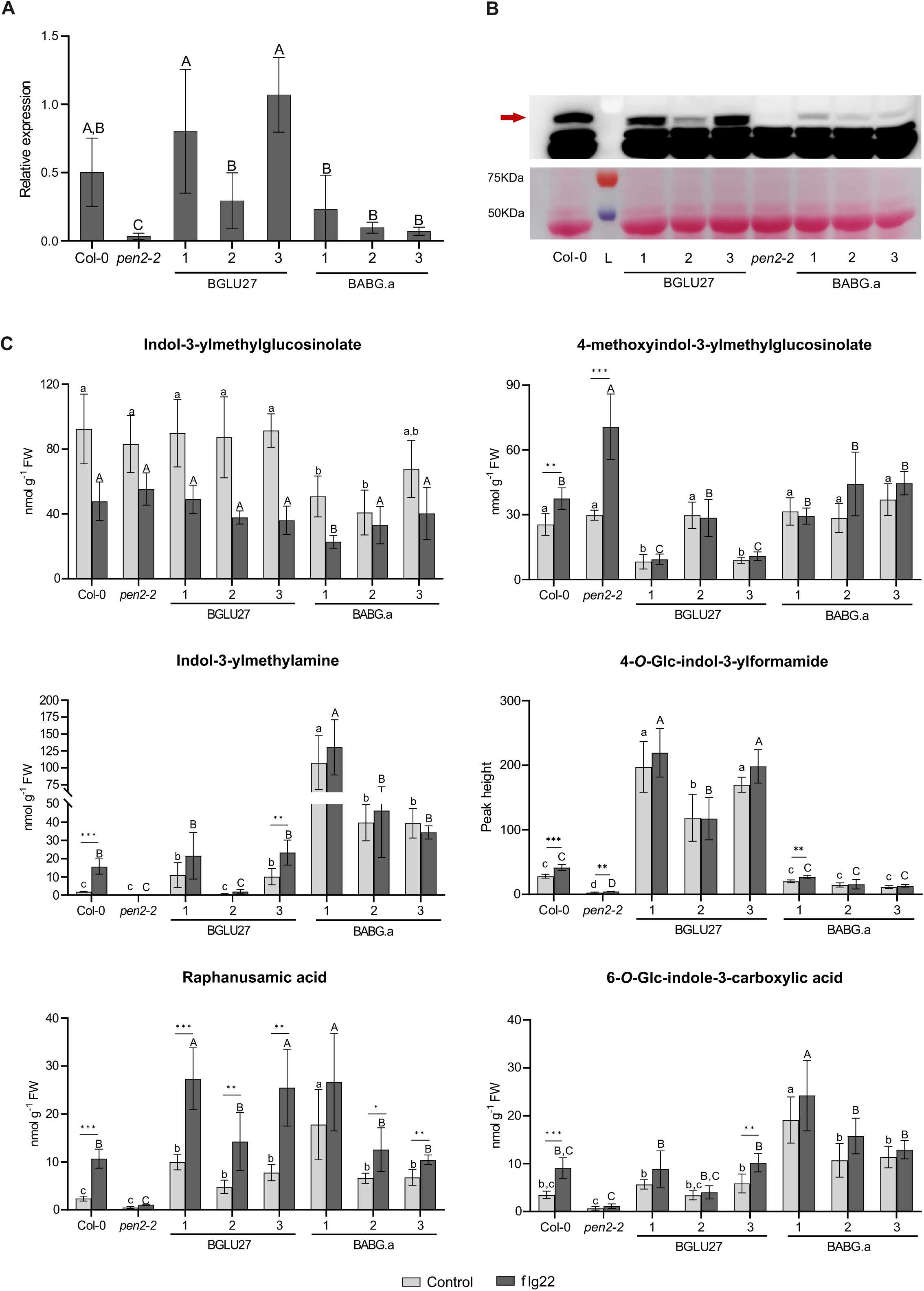
*At*BGLU27 and *Br*BABG.a partially rescue indole glucosinolate hydrolysis in *pen2-2* mutant. **A.** Relative expression of introduced chimeric genes in leaves of selected transgenic lines treated with flg22. Col-0 and *pen2-2* lines are included as positive and negative controls, respectively. **B.** Immunoblot analysis of chimeric BGLUs in leaf extracts from the same set of plant lines. The red arrow indicates position of *At*PEN2 (60 kDa) signal. **C.** Accumulation levels of selected indole glucosinolates and respective hydrolytic products in leaves of control and flg22-treted plants expressing chimeric *AtBGLU27* and *BrBABG.a*. Presented results (**A** & **C**) are means +/− SD. Significantly different statistical groups of genotypes indicated by the analyses of variance (ANOVA; P < 0.05; Tukey’s post-hoc test) are shown with lower case (control) and upper case (flg22) letters. Values marked with asterisks are significantly different from respective controls (Student’s unpaired t-test; * *P* < 0.05, ** *P* < 0.01, *** *P* < 0.001). FW - fresh weight.

Next, we checked if presence of chimeric proteins restores biochemical phenotype of *pen2* mutant plants. PEN2-mediated IGs hydrolysis have been shown to be triggered with pathogen or MAMP recognition (Bednarek *et al*., 2009; Frerigmann *et al*., 2016). Considering this, we analysed accumulation patterns of IG substrates and respective hydrolytic products in leaves of flg22-treated plants to see any potential complementation of PEN2 function by generated chimeric BGLUs in *pen2-2* background. First, we checked accumulation levels of I3G, which does not differentiate Col-0 and *pen2-2* lines. We found a similar accumulation pattern in samples from control and flg22-treated leaves of *At*BGLU27 lines, respectively (Fig. 2C). Interestingly, accumulation levels of this IG were significantly reduced in extracts from control leaves of *Br*BABG.a lines #1 & #2 and from flg22-treated leaves of line #1 as compared with respective Col-0 and *pen2-2* samples. This indicated elevated constitutive I3G hydrolysis in these two lines. This conclusion was further supported with the accumulation levels of I3A (Fig. 2C). This particular metabolite hyperaccumulated constitutively in all *Br*BABG.a lines, reaching the highest accumulation levels in line #1. I3A formation was also restored in *At*BABG27 lines #1 & #3, which showed clearly higher transgene expression and protein accumulation as compared with line #2 (Fig. 2AB). In the case of these two lines, constitutive I3A accumulation levels were higher than in Col-0, however, still much lower than in any of *Br*BABG.a lines. Interestingly, apart from *At*BGLU27 line #3, we did not observe any significant flg22-triggered increase in I3A accumulation in our transgenic plants. This suggested that, unlike in the case of *At*PEN2, I3G hydrolysis driven by *At*BGLU27 and *Br*BABG.a are mainly constitutive, rather than specifically MAMP-triggered.

In case of 4MI3G, the hyperaccumulation observed in *pen2-2* plants was no longer the case in all transgenic lines indicating both chimeric proteins are functional for hydrolysis of 4OH/4MI3G (Fig. 2C). Interestingly, accumulation of 4MI3G was reduced clearly below Col-0 levels in *At*BGLU27 lines #1 & #3. This correlated with accumulation of the corresponding hydrolysis product 4OGlcI3F, which hyperaccumulated in all tested *At*BGLU27 lines. At the same time, accumulation of this compound was restored to Col-0 levels in all *Br*BABGa plants. Overall, this indicated that both enzymes can hydrolyse 4OH/4MI3G, however, *At*BGLU27 is much more reactive towards this substrate(s). Additionally, with the exception of *Br*BABG.a line #1, we did not observe any significant increase in 4OGlcI3F accumulation upon flg22 treatment. This confirmed that unlike *At*PEN2 both enzymes act mainly constitutively. We checked also accumulation of RA, which can be formed from I3G and 4OH/4MI3G hydrolysis in parallel to I3A and 4OGlcI3F (Fig. 1). Interestingly, accumulation pattern of this compound did not represent superposition of trends observed for I3A and 4OGlcI3F (Fig. 2C). Depending on the line and treatment, we observed either Col-0-like or elevated accumulation of RA. Importantly, with the exception of *Br*BABGa line #1, we observed significant induction of RA accumulation upon flg22-tretment. This indicated that apart from the constitutive activity, IG hydrolysis in our lines is still additionally busted with flg22-treatment.

We wondered if the observed constitutive myrosinase activity of chimeric *At*BGLU27 and *Br*BABG.a is their intrinsic property or may occurred as an artefact of our expression system. We used subjectively defined *AtPEN2* promoter region to express our chimeric constructs that did not contain native *PEN2* introns. To check if this could affect spatial or temporal expression of respective transgenes and consequently activity of tested enzymes, we also expressed CDS region of *AtPEN2* under the same promoter region in *pen2-2* plants. After selection of transgenic lines with clearly detectable PEN2 protein (Fig. S4A), we tested them for accumulation of marker metabolites. This revealed predominantly flg22-triggered IG hydrolysis and formation of respective products (Fig. S4B). Particularly metabolic profile of line #3, which possessed Col-0-like PEN2 protein levels, was undisguisable from Col-0. Overall, this indicated that constitutive activity of *At*BGLU27 and *Br*BABG.a is an indigenous property of these myrosinases rather than an artefact of our expression system.

### Chimeric *At*BGLU27 and *Br*BABG.a contribute to immune responses

We next asked the question if our chimeric *At*BGLU27 and *Br*BABG.a proteins can contribute to *A. thaliana* immunity by restricting entry of non-adapted pathogens. To address this question, we inoculated respective transgenic lines together with Col-0 and *pen2-1* plants with the non-adapted on *A. thaliana* hemibiotrophic fungus *Colletotrichum tropicale*. Macroscopic and microscopic inspection of inoculated leaves revealed reduced necrotic lesions on *At*BGLU27 and *Br*BABG.a transgenic lines as compared with *pen2-1* mutants (Fig. 3AB). However, the lesion area on leaves of transgenic plants was still bigger than on the wild type Col-0. We also scored entry of *C. tropicale* into cotyledon epidermal cells of tested lines. Similar as with the lesion areas, entry rates recorded for transgenic lines were significantly reduced as compared with *pen2-1* mutants, but still higher than on Col-0 (Fig. 3C). Overall, these assays indicted that *At*BGLU27 and *Br*BABG.a proteins fused with PEN2-TA can partially replace *At*PEN2 in restricting entry of non-adapted pathogens.

**Figure 3.**
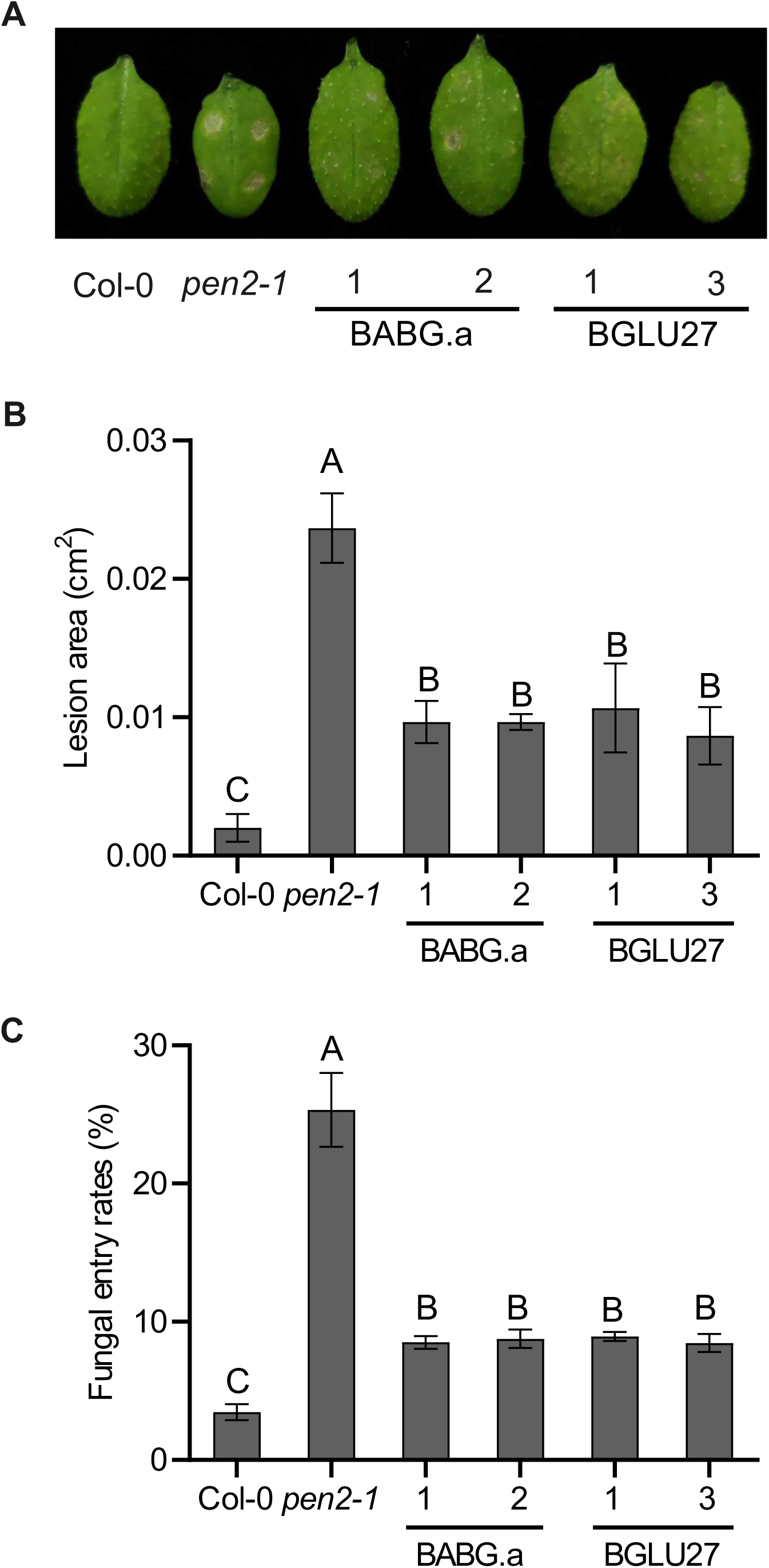
*At*BGLU27 and *Br*BABG.a chimeric proteins partially restore resistance of *pen2-2* plants to *Colletotrichum tropicale*. **A.** Representative leaves of indicated genotypes 4 days after inoculation (dpi) with conidial suspension of *C. tropicale.* **B.** Quantification of lesion development at 5 dpi of conidial suspension of *C. tropicale*. The lesion areas were measured from three independent experiments and for each experiment, a total of 32 drops (spots) were evaluated from at least 6 plants. The statistical significance of differences between means was determined by Tukey’s HSD test. Means not sharing the same letter are significantly different (P < 0.05). **C.** Entry rates of germinated *C. tropicale* conidiospores observed on the indicated genotypes at 14 hours after inoculation, The results are means +/− SD from three independent experiments, where at least 100 germinating conidia were counted in each experiment (n ≥ 300).

Experimental evidence indicated that apart from contributing to the pre-invasive resistance, an unknown product(s) of PEN2 pathway contributes to flg22-triggered biosynthesis of indole-3-carboxylic acid derivatives (ICAs) by controlling expression of respective biosynthetic genes (Frerigmann *et al*., 2016). To check if activity of chimeric *At*BGLU27 and *Br*BABG.a can also promote the synthesis of these indole derivatives, we used HPLC-UV to quantify accumulation levels 6-*O*-Glc-ICA (6OGlcICA) in control and flg22-treated plants of respective transgenic lines. Consistent with our previous report (Frerigmann *et al*., 2016), 6OGlcICA accumulation levels in leaves of flg22-treated *pen2-2* remained unchanged in response to flg22 (Fig. 2C). Expression of *AtBGLU27* or *BrBABG.a* constructs elevated accumulation of this compound in leaves of control and flg22-treated plants, with the only exception of *At*BGLU27 line #2. However, only in *At*BGLU27 line #3 we observed significant increase in accumulation of this metabolite in response to flg22. Interestingly, in leaves of *Br*BABG.a line #1 accumulation levels of 6OGlcICA were higher than in Col-0. Overall, our analysis revealed that similarly like in the case of direct products of PEN2 pathway, expression of both chimeric BGLUs enhances ICA accumulation, however, without clear restoration of increased biosynthesis upon flg22 recognition (Fig. 2C).

### Targeting *At*BGLU18, *At*BGLU23, and *At*BGLU28 to the *At*PEN2 subcellular localization compromises protein stability

Next, we ask the question if more distantly related EE myrosinases can at least partially complement *At*PEN2 function like *At*BGLU27 and *Br*BABG.a. To answer these questions, we selected members of other EE myrosinase subclades including *At*BGLU18 *At*BGLU23/PYK10 and *At*BGLU28 (Fig. S1). Among these, *At*BGLU18 has been suggested to possess myrosinase activity based on metabolic analysis of *bglu18 pyk10* double mutant plants (Nakazaki *et al*., 2019), while myrosinase activities of *At*BGLU23 and *At*BGLU28 have been confirmed in *in vitro* assays (Nakano *et al*., 2017; Sugiyama *et al*., 2021). We generated chimeric CDS similar as for *At*BGLU27 and *Br*BABG.a by trimming sequence encoding N- and C-terminal ER sorting/retention signals from the CDS region of respective genes and added sequence encoding 15aa N-terminal and 65 aa of PEN2-TA (Table 1, Fig. S2, S3A).

After obtaining transgenic lines, we analysed the expression for *AtBGLU28*, *AtBGLU23* and *AtBGLU28* chimeric genes. This indicated that we obtained at least two lines for each chimeric construct with relative transgene expression similar to *AtPEN2* expression in Col-0 (Fig. 4A). However, despite this consistent gene expression, no corresponding protein signals were detected by immunoblot analysis using an antibody specific to the *At*PEN2-TA in any of the tested transgenic lines (Fig. 4B). This suggests probable instability of the chimeric *At*BGLU18, *At*BGLU23, and *At*BGLU28 proteins generated by our expression system. However, we cannot exclude the possibility that degradation of the chimeric proteins is limited to removal of the artificially introduced *At*PEN2-TA, which contains the epitope recognized by the antibody used for western blot analysis. Such limited degradation could still result in active enzymes. To address this possibility, we analysed the accumulation levels of IGs and their hydrolytic products in the respective transgenic lines to detect any potential myrosinase activity associated with *At*BGLU18, *At*BGLU23 or *At*BGLU28. However, no significant changes were observed in the accumulation patterns of the monitored compounds, and all respective transgenic lines exhibited a *pen2-2*-like metabolic phenotype (Fig. S5, S6). Overall, these findings indicate that the tested BGLUs targeted to the subcellular localization of *At*PEN2 are likely degraded by the protein homeostasis machinery of *A. thaliana*.

**Figure 4.**
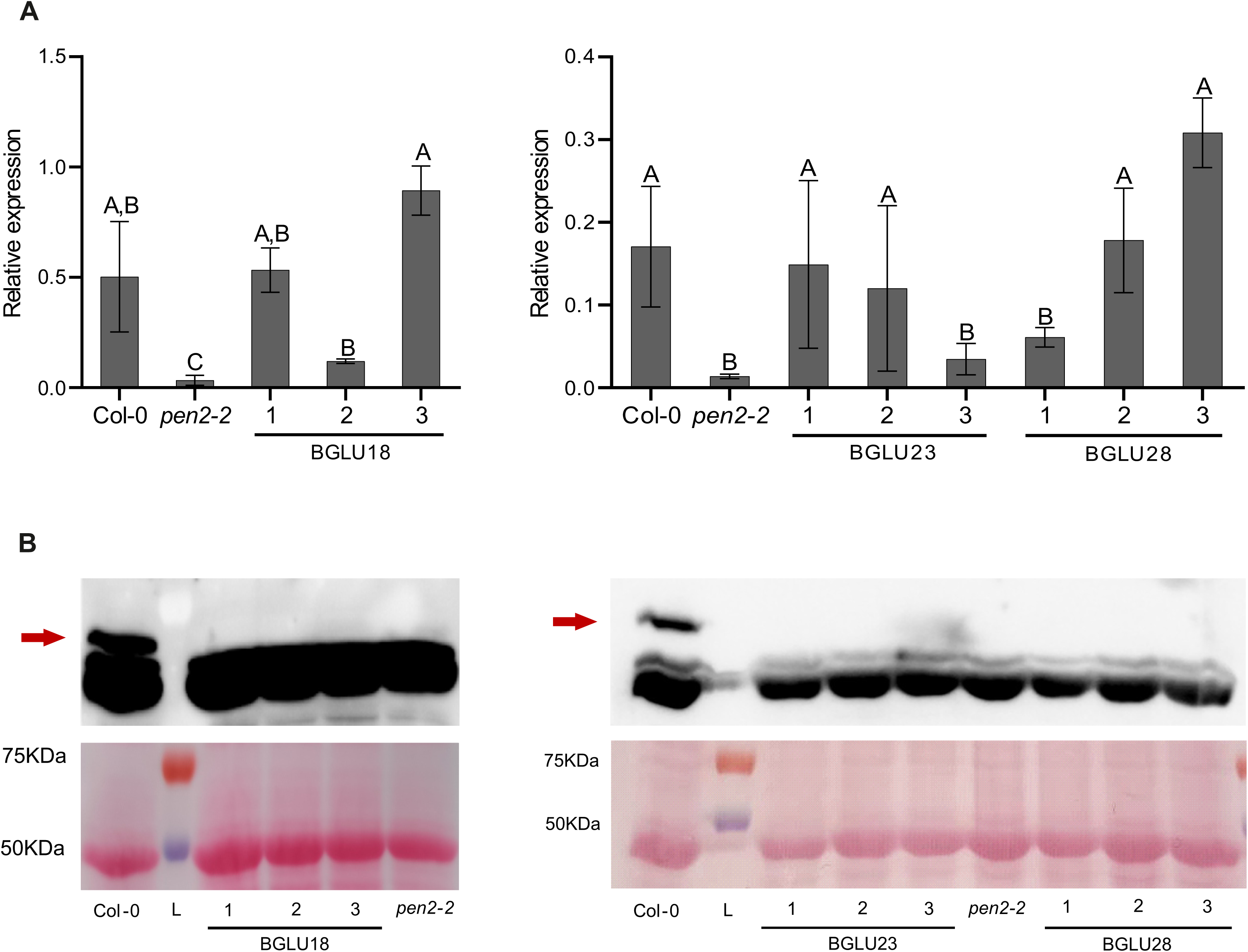
Expression of *AtBGLU18*, *AtBGLU23* and *AtBGLU28* constructs does not result in accumulation of chimeric proteins. **A.** Relative expression of introduced chimeric genes in leaves of selected transgenic lines treated with flg22. Col-0 and *pen2-2* lines are included as positive and negative controls, respectively. Presented results are means +/− SD. Significantly different statistical groups of genotypes indicated by the analyses of variance (ANOVA; P < 0.05; Tukey’s post-hoc test) are shown with upper case letters. **B.** Immunoblot analysis of chimeric BGLUs in leaf extracts from the same set of plant lines. The red arrow indicates position of *At*PEN2 (60 kDa) signal.

### *At*PEN2*, At*BGLU27 and *Br*BABG.a lack conserved post-translational modifications

Next, we asked the question about molecular determinants of instability of chimeric *At*BGLU18, *At*BGLU23 and *At*BGLU28 proteins. To answer this, we delved into the protein sequences of all 47 *At*BGLUs and two *Br*BABGs from *B. rapa.* We noted that the vast majority of *A. thaliana* BGLU family members, with the exception of *At*PEN2, *At*BGLU27, and *At*BGLU42, are predicted to contain N-terminal signal peptides (Xu *et al*., 2004). Moreover, based on the UniProt database entries (UniProt Consortium, 2025), we identified conserved Asn and Cys residues predicted to be involved in post-translational modifications in most *At*BGLUs (Fig. 5A, Table 1). These modifications included *N*-glycosylations and disulfide bonds (DSBs), respectively. *N*-glycosylation of respective Asn residues has been confirmed by proteomic analysis of *At*TGG1 and *At*TGG2 (Liebminger *et al*., 2012). The DSB formation by the two strongly conserved Cys residues (Fig. 5A) has been observed in crystal structures of a cyanogenic BGLU from white clover (Barrett *et al*., 1995), BGLU from maize (Zouhar *et al*., 2001) as well as in *Os*3BGLU6 and *Os*4BGLU18 from rice (Seshadri *et al*., 2009; Baiya *et al*., 2021). The number and location of putative *N*-glycosylation sites were found to be variable in *At*BGLUs (Table 1). Notably, all of the conserved Asn residues were exclusively absent in *At*BGLU25-27 and *Br*BABGs. In addition, we found the two Cys residues forming the conserved DSB not fully retained in *At*PEN2, *At*BGLU27 and *Br*BABG.a/b (Fig. 5A, Table 1). *At*BGLU26 lacks one of these, while *At*BGLU27 and *Br*BABGs lacks both Cys residues. Among other *At*BGLUs, the second Cys was absent in *At*BGLU7 and *At*BGLU8, while both Cys residues are absent from *At*BGLU39 and *At*BGLU42 (Fig. 5A & Table 1). However, *AtBGLU39* was suggested to be a pseudogene because of the presence of few frameshift mutations (Xu *et al*., 2004; Wang *et al*., 2009). Overall, we found that *At*PEN2, *At*BGLU27 and *Br*BABGs differ from other *At*BGLUs by their deficiency in N-terminal targeting signals and lack of all post-translational modifications.

**Figure 5.**
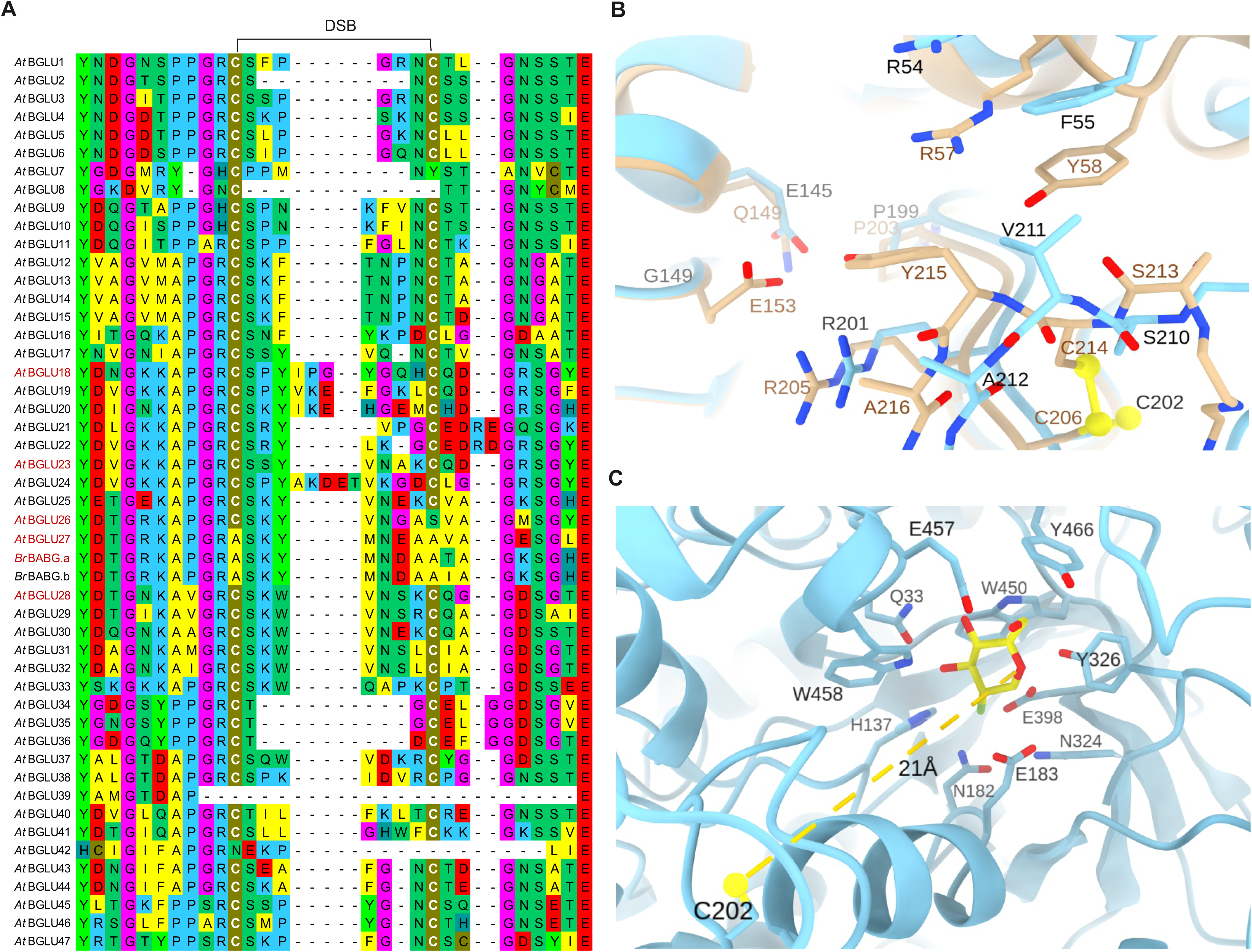
The strongly conserved disulfide bond (DSB) is absent in *At*PEN2, *At*BGLU27 and *Br*BABGs. **A.** Multiple sequence alignment of *At*BGLU and *Br*BABG regions containing two conserved Cys residues involved in the DSB formation. BGLUs investigated in this study are highlighted in red. **B.** Structural comparison of *At*PEN2/BGLU26 and *Sinapis alba* myrosinase. The AlphaFold model of *At*PEN2 is shown in light blue, and the experimentally determined structure of *S. alba* myrosinase (PDB ID: 1MYR) is shown in khaki. Residues located within a 4 Å radius are labelled in black for *At*PEN2 and brown for *S. alba* myrosinase. Sulfur atoms are shown as yellow spheres. **C.** Location of C^202^ relative to the active site in the AlphaFold model of *At*PEN2. The active site was mapped by superposition with the 1E70 structure; the G2F ligand from 1E70 is shown in yellow. The distance to C^202^ is indicated by a yellow dashed line. Residues within 4 Å from the superposed ligand are shown and labelled. E^183^ and E^398^ that correspond to Q^187^ and E^409^ in *S. alba* myrosinase, respectively.

### Restoration of DSB in *At*PEN2 disrupts its activity

Next, we put efforts to understand the possible role of the observed loss of these post-translational modifications, particularly selective absence of Cys residues predicted to be involved in DSB in *At*PEN2, *At*BGLU27 and *Br*BABGs. To evaluate the phenotypic and metabolic consequences of DSB formation in *At*PEN2, we generated transgenic *pen2-2* plants expressing mutated variant of this enzyme that was designed to reintroduce DSB (PEN2_DSB). Inspection of the ^210^SVA^212^ region in *At*PEN2, aligned with the corresponding ^213^SCYA^216^ segment of *S. alba* myrosinase, indicated that restoring the putative DSB geometry would require extension of this loop by one residue (Fig. 5B). In the *S. alba* structure, Tyr^215^ occupies a niche formed by residues that are conserved or similar in *At*PEN2, including Arg^201^, Pro^199^, and Glu^145^, which correspond to Arg^205^, Pro^203^, and Gln^149^ in *S. alba* myrosinase, respectively. Therefore, Val^211^ in *At*PEN2 was replaced with a Cys-Tyr dipeptide (Fig. S7A). This substitution was expected to generate a local arrangement compatible with the formation of a DSB, while the introduced Tyr residue was expected to preserve the aromatic side chain packing observed in the reference structure and thereby support the geometry required for DSB formation and/or its stabilization (Fig. 5B).

To assess the effect of the introduced substitution, firstly, we analysed the expression of *PEN2_DSB* in selected transgenic lines. We found the transgene expression similar to *AtPEN2* in Col-0 (Fig. 6A). The gene expression was also complemented by consistent protein signals in PEN2_DSB lines corresponding to *At*PEN2 in Col-0 in immunoblot analysis (Fig. 6B). Next, we asked the question if the mutation indeed resulted in DSB formation. In eukaryotic cells, stable structural DSBs are formed mainly in proteins that enter the secretory pathway, because the ER lumen is an oxidizing folding environment and contains dedicated enzymes, including disulfide isomerase (Tu & Weissman 2004). *At*PEN2 as a protein with a C-terminal membrane TA is not predicted to enter the secretory pathway and remains cytosolic (Marty *et al*., 2014; Fuchs *et al*., 2015). Unlike ER, the cytosol is generally a reducing compartment, therefore stable DSBs are uncommon in cytosolic proteins. To validate DSB formation in PEN2_DSB, we pre-fractionated protein extracts from leaves of Col-0 and PEN2_DSB transgenic lines using electrophoresis in non-reducing conditions. We digested fractions containing PEN2/PEN2_DSB proteins with trypsin and subjected them to LC-MS analysis. Tryptic digestion of the DSB region of PEN2 should release two peptides: ^202^CSK^204^ (Pep1) and ^205^YVNGAS**<u>V</u>**AGMSGYEAYIVSHNMLLAHAEAVEVFR^238^ (Pep2). The mutation introduced in PEN2_DSB changes Pep2 into ^205^YVNGAS**<u>CY</u>**A…R^239^ (Pep3). In case of successful DSB formation, Pep1-Pep3 adduct should be released after the tryptic digestion. During our LC-MS analysis, we detected signal corresponding to Pep2 in samples from Col-0, but not from PEN2_DSB lines (Fig. S7C). Conversely, a signal matching with Pep1-Pep3 adduct was exclusively present in samples from PEN2-DSB lines (Fig. S7C), revealing that our mutation led to efficient DSB formation. During the same analysis, we did not detect any signals that could represent single Pep3. This indicated that the PEN2_DSB variant without formed DSB is below our detection limit. Next, we tested the impact of the introduced DSB on PEN2 function. Our targeted HPLC-UV/FLD analysis revealed *pen2*-like accumulation profile of IGs and respective hydrolytic products in leaves of control and flg22-treated PEN2_DSB plants (Fig. 6C). This indicated that presence of DSB led to loss of hydrolytic activity of *At*PEN2.

**Figure 6.**
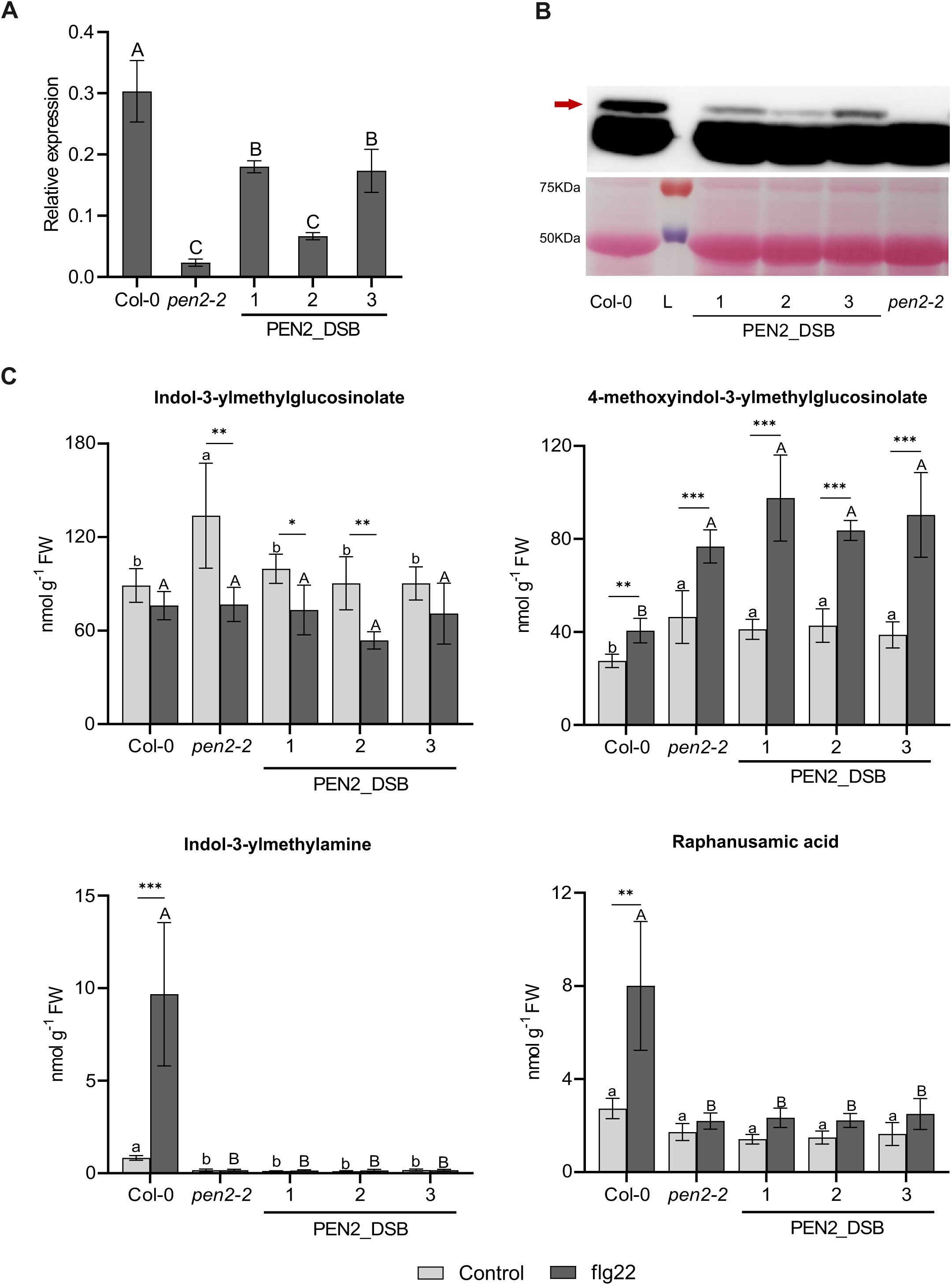
Introduction of a disulfide bond (DSB) abolish *At*PEN2 activity. **A.** Relative expression of mutated *AtPEN2* gene (*PEN2_DSB*) in leaves of selected transgenic lines treated with flg22. Col-0 and *pen2-2* lines are included as positive and negative controls, respectively. **B.** Immunoblot analysis of PEN2_DSB protein in leaf extracts from the same set of plant lines. The red arrow indicates position of *At*PEN2 (60 kDa) signal. **C.** Accumulation levels of selected indole glucosinolates and respective hydrolytic products in leaves of control and flg22-treated plants expressing *PEN2_DSB* construct. Presented results (**A** & **C**) are means +/− SD. Significantly different statistical groups of genotypes indicated by the analyses of variance (ANOVA; P < 0.05; Tukey’s post-hoc test) are shown with lower case (control) and upper case (flg22) letters. Values marked with asterisks are significantly different from respective controls (Student’s unpaired t-test; * *P* < 0.05, ** *P* < 0.01, *** *P* < 0.001). FW - fresh weight.

### PEN2, BGLU27 and BABG proteins are highly conserved in Brassicaceae

Our analysis revealed that *At*PEN2, *At*BGLU27, and *Br*BABG differ from other myrosinases and from the majority of BGLUs in lacking post-translational modifications (Table 1; Fig. 3A). To learn more about conservation of these unique enzymes within Brassicaceae family we searched available genomic data at Phytozome and other databases for homologs of these proteins (Goodstein *et al*., 2012; Kagale *et al*., 2014; Willing *et al*., 2015; Gan *et al*., 2016; Nguyen *et al*., 2019). We also included in this search available genomes of species representing other families belonging to Brassicales order and known to produce IGs (Edger *et al*., 2018). These included *Cleome spinosa*, *Tarenaya hassleriana* and *Gynandropsis gynandra* (Cleomaceae) as well as *Capparis spinosa* (Capparaceae) (Goodstein *et al*., 2012; Cheng *et al*., 2013; Wang *et al*., 2022; Hoang *et al*., 2023). Next, we performed phylogenetic analysis of the identified sequences (Fig. 7, Table S2). This revealed presence of PEN2 orthologs in majority of the analysed Brassicaceae species except *Capsella* spp. and *Camelina sativa*, which have been characterized to lose the capacity to produce IGs (Czerniawski *et al*., 2021). In addition, PEN2 orthologs were also present in species representing Cleomaceae and Capparaceae families indicating this unique myrosinase evolved before the speciation of Brassicaceae, Cleomaceae and Capparaceae.

**Figure 7.**
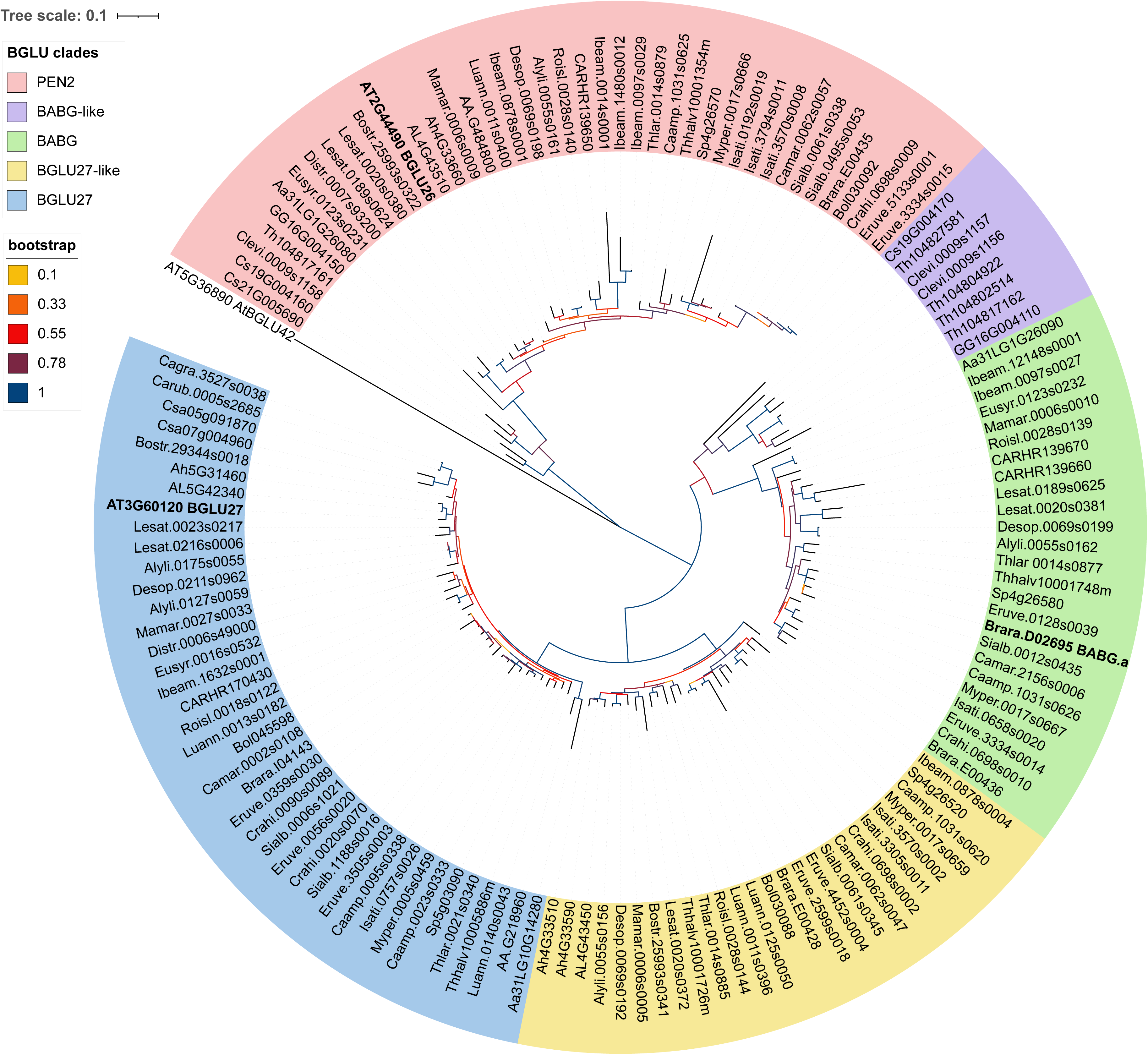
PEN2, BABG/BABG-like and BGLU27/BGLU27-like proteins are conserved within Brassicaceae, Cleomaceae, and Capparaceae families. The maximum-likelihood phylogenetic tree of putative orthologs and close homologs of investigated BGLUs identified in available genomes of Brassicaceae, Cleomaceae, and Capparaceae species. For species abbreviations see Table S2.

Our analysis revealed the presence of *Br*BABG orthologs in most lineages of the Brassicaceae family. Notably, these orthologs were completely absent from the Arabidopsideae, Boechereae, and Camelineae tribes, indicating that BABG was likely lost in the common ancestor of these lineages (German *et al*., 2023)(Fig. 7, Table S2). We also identified highly homologous BGLUs in *C. violacea*, *T. hassleriana*, *G. gynandra* and *C. spinosa*, however, their sequences formed a distinct sub-clade that looked separated from the core BABG clade in the phylogenetic tree. We therefore referred to these sequences as BABG-like.

Finally, we found *At*BGLU27 orthologs in all analysed Brassicaceae species. In addition, we identified a new homolog of this enzyme, which we named BGLU27-like (Fig. 7, Table S2). Although BGLU27-like is absent from the reference *A. thaliana* genome, orthologous genes were present in the vast majority of analysed Brassicaceae genomes, including those of *Arabidopsis lyrata* and *Arabidopsis halleri*. In contrast to PEN2 and BABG/BABG-like, we did not identify any putative *At*BGLU27 or BGLU27-like orthologs in the investigated Cleomaceae and Capparaceae species. This suggests that BGLU27 and BGLU27-like are evolutionarily more recent than PEN2 and BABG/BABG-like and evolved after the divergence of the Brassicaceae lineage from the ancestors of Cleomaceae and Capparaceae (Edger *et al*., 2018).

In addition, to the phylogenetic analysis, we also checked the conservation of the investigated DSB in the identified orthologs. We found the second Cys is missing in all identified *At*PEN2 orthologs, while all *At*BGLU27, BGLU27-like and *Br*BABG orthologs lack both Cys residues (Fig. S8, S9). Interestingly, the two *At*PEN2 orthologs identified in *C. spinosa* also lacked both Cys residues. Finally, the identified BABG-like proteins lack only the first Cys residue, confirming that these enzymes are somehow distinct from BABGs (Fig. S9).

## Discussion

### PEN2-related myrosinases differ in pathogen-triggered activation and substrate preference

In this study, we addressed the molecular determinants of the *At*PEN2 function in pathogen-triggered IG-metabolism. We found that the two closest homologs, *At*BGLU27 and *Br*BABG.a can partially complement *At*PEN2 function when anchored in the mitochondrial membrane via PEN2-TA. This included hydrolysis of IGs to I3A, 4OGlcI3F and RA as well as control of entry of non-adapted pathogens (Fig. 2, Fig. 3). However, changes in the accumulation levels of IGs and their hydrolytic products observed in transgenic lines indicated that even targeted to the same intracellular compartment, *At*BGLU27 and *Br*BABG.a differ from *At*PEN2 being constitutively active. This suggests the existence of unique molecular mechanisms that activate *At*PEN2 exclusively upon pathogen recognition. In this context, it is noteworthy that, according to *in vitro* assays with recombinant protein, this BGLU is active only over a relatively narrow pH range, with an optimum around pH 6.0 (Bednarek *et al*., 2009). This contrasts, for instance, with *At*TGG1 and *At*TGG2, which remain active across a broad pH range, from pH 4.5 to 10 (Zhou *et al*., 2012). Moreover, although *At*PEN2 is anchored in the mitochondrial membrane, the protein is exposed to the cytoplasm, where the typical pH of 7.2-7.5 does not correspond to the range of PEN2 activity (Felle, 2001; Bednarek *et al*., 2009). This mismatch could cause the constitutive inactivity of this BGLU under resting cellular conditions. Ion fluxes and redox processes at the cell periphery, beneath attempted pathogen entry sites where *At*PEN2 re-localizes, may lead to local cytoplasmic acidification and thereby trigger *At*PEN2 activity (Fuchs *et al*., 2015; Kesten *et al*., 2019; Li *et al*., 2025). Despite its molecular basis, such activation mechanism prevents from constitutive formation of potentially toxic end products and from the reduction of defensive potential through depletion of the native glucosinolate pool. This mechanism appears to be absent from *At*BGLU27 and *Br*BABG.a, which, based on our experiments, are active under native cytoplasmic pH conditions.

### *At*BGLU27 and *Br*BABG.a reveal partial functional overlap with PEN2-dependent immunity

Despite constitutive activity of the chimeric *At*BGLU27 and *Br*BABG.a proteins, analysis of RA accumulation levels in control and flg22-treated leaves indicated that both enzymes are able to increase IG hydrolysis upon re-localization to the pathogen contact sites. However, this increase was not evident from the accumulation of I3A and 4OGlcI3F (Fig. 2). This discrepancy probably results from the reported retrograde metabolism of RA to Cys (Sugiyama *et al*., 2021), which likely efficiently removes the excess of constitutively generated RA making the flg22-triggered changes apparent. In contrast, I3A and 4OGlcI3F are likely less readily, if at all, metabolized and therefore overaccumulate *in planta* obscuring the impact of flg22-induced IG hydrolysis on their formation. The pathogen triggered IG-hydrolysis have been additionally confirmed with partial rescue of the infection phenotypes. Both proteins significantly reduced entry rates of *C. tropicale* as compared with the *pen2-1* mutant background (Fig. 3). The reduced entry rates corresponded with reduced size of the lesions observed at the droplet inoculation sites. However, entry rates and lesion size have been not reverted to the Col-0 levels. This partial rescue of the immune response could result from lower capacity of *At*BGLU27 and *Br*BABG.a to hydrolyse IGs at the sites of attempted entry as compared with *At*PEN2, or with reduced availability of IGs due to the observed constitutive activity of both enzymes. However, IG biosynthesis, including CYP81F2 mediated modification of the indole core, have been reported to be induced specifically in the pathogen-inoculated cells, so the latter option seems to be less likely (Fuchs *et al*., 2015; Hunziker *et al*., 2020).

Interestingly, even though both investigated BGLUs were able to partially replace *At*PEN2, they exhibited different substrate specificity towards unsubstituted and substituted IGLs. *Br*BABG.a lines accumulated high levels of I3A and reduced amounts of I3G indicting this enzyme efficiently hydrolyse I3G (Fig. 2). Conversely, lines expressing *At*BGLU27 hyper-accumulated 4OGlcI3F and were depleted in 4MI3G revelling BGLU27 prefers this substituted IGL as its substrate. Such substrate specificity is not common among myrosinases. For instance, according to *in vitro* assays, *At*TGG1 and *A*tTGG2 hydrolyse efficiently all types of GLs. *At*BGLU28 has been shown to be more selective and hydrolyse efficiently few selected GL individuals (Sugiyama *et al*., 2021). This observed striking difference in substrate specificity between *At*BGLU27 and *Br*BABG.a can be linked with functions of these enzymes. *Br*BABGs have been shown to be require for brassinin biosynthesis, which starts with I3G hydrolysis (Klein & Sattely, 2017). Therefore, specificity towards this substrate seems reasonable. On the other hand, analysis of our transgenic lines indicted that despite strong preference towards I3G this enzyme can still hydrolyze 4MI3G, which matches with the reported 4-methoxybrassinin accumulation in *Brassica oleracea* (Monde *et al*., 1990).

Unlike *Br*BABG.a, the function of *At*BGLU27 remains obscure. Based on published results, this enzyme appears to be stress- and pathogen-responsive. *AtBGLU27* expression has been reported to be strongly induced by cellobiose in *A. thaliana* seedlings and by the pathogenic fungus *Verticillium longisporum* in roots (Iven *et al*., 2012; Souza *et al*., 2017). In the latter case, transcriptional co-induction of *CYP81F2* and other tryptophan metabolism-related genes was observed, suggesting that this BGLU may be involved in pathogen-triggered IG metabolism (Iven *et al*., 2012). Our results are consistent with these findings, as they reveal myrosinase activity of *At*BGLU27 pointing to 4OHI3G and/or 4MI3G as physiologically relevant substrate(s) of this enzyme. This substrate specificity is also consistent with the reported co-expression of *AtBGLU27* and *CYP81F2* (Iven *et al*., 2012). Overall, these results suggest that *At*BGLU27 acts as a myrosinase involved in *A. thaliana* immunity by hydrolysing IGs modified by the CYP81F2 monooxygenase. However, *bglu27* mutant plants did not show significantly altered disease symptoms during colonization by *V. longisporum*, indicating that *At*BGLU27 is either functionally redundant, condition-specific, or contributes only weakly to measurable resistance under the tested conditions (Iven *et al*., 2012).

### Secretory-pathway dependence limits functional replacement by other BGLUs

In contrast to *At*BGLU27 and *Br*BABG.a, our attempts to express *At*BGLU18, *At*BGLU23, and *At*BGLU28 in the *At*PEN2-like setup were unsuccessful. Although transcript accumulation was clearly confirmed, the corresponding proteins could not be detected by western blotting (Fig. 4B). This suggests that the respective chimeric enzymes were unstable. In this context, we noted that plant BGLUs are often routed through the secretory pathway. Correspondingly, most of *A. thalinana* family members, except *At*PEN2, *At*BGLU27, and *At*BGLU42, are predicted to contain N-terminal signal peptides and consequently to localize to the ER, vacuole, apoplast, or related compartments (Xu *et al*., 2004). For instance, the whole *At*BGLU18-24 clade possess predicted N-terminal signal peptides as well as ER-retention signals at their C-termini (Table 1) (Nagano *et al*., 2008; Ogasawara *et al*., 2009; Nakazaki *et al*., 2019). Moreover, imaging of lines expressing fluorophore tagged proteins revealed that *At*BGLU18 and *At*BGLU23, the two enzymes tested in our study, localize to ER-bodies, which are unique ER-derived compartments in Brassicaceae (Matsushima *et al*., 2003; Ogasawara *et al*., 2009; Yamada *et al*., 2020). The subcellular localization of *At*BGLU28 has not, to our knowledge, been experimentally resolved. Studies on sulfur-deficiency-induced glucosinolate catabolism discuss *At*BGLU28 as a putative vacuolar enzyme (Zhang *et al*., 2020). This proposed localization would be consistent with a role in hydrolysing vacuole stored glucosinolates in response to sulfur-starvation, but direct experimental validation is still lacking.

Regardless of their final localization, all BGLUs routed through the secretory pathway are expected to undergo post-translational modifications, including *N*-glycosylation and DSB formation. This is supported by the presence of conserved Asn and Cys residues in these proteins (Table 1). In the case of myrosinases, *N*-glycosylation has been confirmed by crystal-structure studies of the *S. alba* myrosinase and by proteomic analyses of *At*TGG1 and *At*TGG2 (Burmeister *et al*., 1997; Liebminger *et al*., 2012). DSB formation has not been directly confirmed for any myrosinase or *At*BGLU, but it has been unambiguously demonstrated in several BGLUs from other species, including white clover, maize and rice (Barrett *et al*., 1995; Zouhar *et al*., 2001; Seshadri *et al*., 2009; Baiya *et al*., 2021). The strong conservation of the corresponding Cys residues in most *At*BGLUs suggests that this particular DSB may also form in these proteins (Table 1).

As mentioned above, in contrast to the majority of *At*BGLUs, *At*PEN2 and *At*BGLU27 do not contain predicted N-terminal signal peptides (Xu *et al*., 2004). Moreover, as a protein with a C-terminal TA, *At*PEN2 is not predicted to enter the secretory pathway but instead remains cytosolic (Marty *et al*., 2014; Fuchs *et al*., 2015). Consequently, the designed chimeric proteins, depleted of their native N- and C-terminal peptides and fused to the *At*PEN2-TA, are unlikely to pass through the secretory pathway. They are therefore not expected to undergo post-translational modifications that may be crucial for proper folding. This could lead to protein instability and may explain the absence of detectable signals in the western blot analysis.

### DSB loss enabled cytosolic PEN2-related myrosinase activity

The lack of predicted signal peptides in *At*PEN2 and *At*BGLU27 correlates with the loss of conserved Asn and Cys residues potentially involved in post-translational modifications (Table 1). Similar features are also observed in *Br*BABGs, which are close homologs of *At*PEN2 and *At*BGLU27. In addition to these enzymes, *At*BGLU25 has lost all conserved Asn residues; however, this protein retains both conserved Cys residues and has been shown to localize to the ER (Cho *et al*., 2025). Conversely, the Cys residues potentially involved in DSB formation are not conserved in *At*BGLU7, *At*BGLU8, and *At*BGLU42. Among these proteins, *At*BGLU42 has been suggested to be cytoplasmic based on the absence of an obvious N-terminal signal peptide (Xu *et al*., 2004; Horikoshi *et al*., 2022). If this is the case, the conserved Asn residue in *At*BGLU42 would no longer be *N*-glycosylated and, similarly to *At*PEN2 and *At*BGLU27, this enzyme would be expected to lack all post-translational modifications.

According to our genomic and phylogenetic analyses, the loss of post-translational modification sites, particularly those associated with DSB formation, is conserved in all putative orthologs and close homologs of *At*PEN2, *At*BGLU27, and *Br*BABG, indicating that this loss had already occurred in the common ancestor of these enzymes. (Fig. 6, Fig. S8, Fig. S9). Strikingly, targeted mutagenesis of the *At*PEN2 sequence indicated that the DSB loss is indispensable for *At*PEN2 activity. These results suggest that the DSB formation may act as a regulatory element controlling BGLU activity, with its presence maintaining the enzyme in an inactive state. In cytoplasmic enzymes, loss of this DSB may therefore represent an adaptation enabling activity outside the secretory pathway. However, the position of this DSB within the protein structure does not readily support such a mechanism. The DSB is located at the entrance to the active-site cavity, but far from the catalytic center. Specifically, the distance from Sγ of Cys^202^ in BGLU26 to the centroid of the 2-deoxy-2-fluoro-α-D-glucopyranose ligand (G2F) from the superposed 1E70 structure is approximately 21 Å (Fig. 5C) (Burmeister *et al*., 1997). In any case, such a control function seems likely as DSB formation has been shown to affect activity of different type of enzymes, including a member of glycosyl hydrolase family 2 (Martins *et al*., 2026).

### Evolution of PEN2-related myrosinases parallels Brassicales immune chemistry

In addition to revealing the conserved absence of the DSB, our phylogenetic analysis showed strong conservation of PEN2, BGLU27, BGLU27-like, and BABG proteins within the Brassicaceae family. The only exceptions were the loss of *At*PEN2 orthologs in the Camelineae I tribe and the loss of *Br*BABG orthologs in the Arabidopsideae, Boechereae and Camelineae I tribes. The loss of PEN2 has been shown to correlate with the loss of IG biosynthetic capacity in the respective species (Bednarek *et al*., 2011; Czerniawski *et al*., 2021). In contrast, the absence of *Br*BABG orthologs correlates with differences in the occurrence and biosynthesis of Brassicaceae phytoalexins (Pedras *et al*., 2011). Members of the Arabidopsideae and Camelineae I tribes produce camalexin as their major phytoalexin (Bednarek *et al*., 2011). Unlike brassinins and some other phytoalexins, such as wasalexins, camalexin is not derived from IGs, rendering BABG dispensable in these lineages (Glawischnig, 2007; Pedras *et al*., 2010; Klein & Sattely, 2017).

In addition to being conserved in Brassicaceae, PEN2 orthologs were also identified in Cleomaceae and Capparaceae species. In the same species, we also identified BGLUs highly homologous to BABG, which we refer to as BABG-like. Notably, some *Capparis* spp. have been reported to accumulate sulfur-containing indole compounds, referred to as capparine-like alkaloids, which are structurally analogous to spirobrassinin (Li *et al*., 2008; Zhou *et al*., 2010; Ba *et al*., 2025). These findings indicate that such compounds may be produced from IGs with the contribution of BABG-like enzymes. Unlike PEN2 and BABG/BABG-like, BGLU27 and BGLU27-like do not have detectable orthologs in the investigated genomes of Cleomaceae or Capparaceae species, suggesting that these enzymes are evolutionarily more recent than PEN2 and BABG/BABG-like.

## Conclusions

Overall, our findings indicate that PEN2-related myrosinases represent a Brassicaceae immune-associated BGLU lineage that has diverged from secretory-pathway-dependent EE-myrosinases. *At*BGLU27 and *Br*BABG.a can partially substitute for *At*PEN2 when redirected to its subcellular context, but their constitutive activity and distinct substrate preferences show that *At*PEN2 possesses additional regulatory features required for pathogen-triggered glucosinolate metabolism. The conserved loss of the DSB in PEN2, BGLU27 and BABG proteins, together with the loss of *At*PEN2 activity after DSB restoration, suggests that removal of this structural element was a key evolutionary step enabling cytosol-compatible immune myrosinase function. Thus, PEN2 immune activity appears to result from the combination of subcellular retargeting, loss of secretory-pathway-associated structural constraints and acquisition of regulatory features that restrict glucosinolate hydrolysis to the sites of attempted fungal penetration.

## Acknowledgements

This work was supported by the National Science Centre OPUS grant (UMO-2018/31/B/NZ3/03626) to PB, and Grant-in-Aid for Scientific Research (23K20042 and 25H00431) to YT. We also acknowledge the use of the infrastructure developed under the project NEBI - National Research Center for Imaging in the Biological and Biomedical Sciences, POIR.04.02.00-00-C004/19, co-financed through the European Regional Development Fund (ERDF) in the frame of Smart Growth Operational Programme 2014-2020 (Measure 4.2 Development of modern research infrastructure of the science sector).

## Author contributions

GS, HA, MR, YT and PB planned and designed the research. GS, HA, MPB, SSO, CJ, AP, MB, SK and AS performed experiments, collected and analysed the data. GS, HA, LM, MR, YT and PB interpreted the data and wrote the manuscript. GS and HA contributed equally.

## Supplementary figure legends

**Figure S1 QE- and EE-myrosinases in *Arabidopsis thaliana*.** The maximum-likelihood phylogenetic tree of *At*BGLUs proposed to function as myrosinases. The positions of highlighted conserved catalytic (E/Q) and substrate recognition (RK) residues are based on Nakano *et al*. (2017).

**Figure S2 Conservation of BGLUs selected for this study.** Protein sequence alignment of *At*BGLU26/PEN2 (AT2G44490), *At*BGLU18 (AT1G52400), *At*BGLU27 (AT3G60120), *Br*BABG.a (Brara.D02695), *At*BGLU23/PYK10 (AT3G09260) and *At*BGLU28 (AT2G44460) displaying conserved and variable regions among these myrosinases. The unique 15 aa N-terminal and 65 aa C-terminal regions of *At*PEN2 are underlined with blue and yellow colours, respectively. The C-terminal regions confining ER retention signal in *At*BGLU18 and *At*BGLU23 are highlighted in green colour.

**Figure S3. Design of constructs used in this study. A**. Simplified exemplary map of the T-DNA fragment of the vector used for expression of *At*PEN2 under the control of its native promoter and for the preparation of the remining plasmids. **B**. Chimeric genes/CDS designed for expression of *AtBGLU26/PEN2* (AT2G44490), *AtBGLU18* (AT1G52400), *AtBGLU27* (AT3G60120), *BrBABG.a* (Brara.D02695), *AtBGLU23/PYK10* (AT3G09260) and *AtBGLU28* (AT2G44460). Sequences encoding unique PEN2 N- and C-terminal extensions are indicated in blue and yellow, respectively.

**Figure S4. Expression of *AtPEN2* in *pen2-2* mutant fully restores metabolic phenotype. A.** Immunoblot analysis of *At*PEN2 protein in leaf extracts from selected transgenic lines treated with flg22. Col-0 and *pen2-2* lines are included as positive and negative controls, respectively. The red arrow indicates position of *At*PEN2 (60 kDa) signal. **B.** Accumulation levels of selected indole glucosinolates and respective hydrolytic products in leaves of control and flg22-treated plants expressing *AtPEN2* construct. Presented results are means +/− SD. Significantly different statistical groups of genotypes indicated by the analyses of variance (ANOVA; P < 0.05; Tukey’s post-hoc test) are shown with lower case (control) and upper case (flg22) letters. Values marked with asterisks are significantly different from respective controls (Student’s unpaired t-test; * *P* < 0.05, ** *P* < 0.01, *** *P* < 0.001). FW - fresh weight.

**Figure S5. Expression of *AtBGLU18* construct does not alter accumulation of indole glucosinolates and related hydrolytic products.** Accumulation levels of selected indole glucosinolates and respective hydrolytic products in leaves of control and flg22-treted plants expressing chimeric *AtBGLU18*. Col-0 and *pen2-2* lines are included as positive and negative controls, respectively. Presented results are means +/− SD. Significantly different statistical groups of genotypes indicated by the analyses of variance (ANOVA; P < 0.05; Tukey’s post-hoc test) are shown with lower case (control) and upper case (flg22) letters. Values marked with asterisks are significantly different from respective controls (Student’s unpaired t-test; * *P* < 0.05, ** *P* < 0.01, *** *P* < 0.001). FW - fresh weight.

**Figure S6. Expression of *AtBGLU23* and *AtBGLU28* constructs does not alter accumulation of indole glucosinolates and related hydrolytic products.** Accumulation levels of selected indole glucosinolates and respective hydrolytic products in leaves of control and flg22-treted plants expressing chimeric *AtBGLU23 and AtBGLU28*. Col-0 and *pen2-2* lines are included as positive and negative controls, respectively. Presented results are means +/− SD. Significantly different statistical groups of genotypes indicated by the analyses of variance (ANOVA; P < 0.05; Tukey’s post-hoc test) are shown with lower case (control) and upper case (flg22) letters. Values marked with asterisks are significantly different from respective controls (Student’s unpaired t-test; * *P* < 0.05, ** *P* < 0.01, *** *P* < 0.001). FW - fresh weight.

**Figure S7. Disulfide bond (DSB) is successfully restored in mutated *At*PEN2 protein (PEN2_DSB). A.** Sanger sequencing of the mutated *AtPEN2* CDS to validate V^211^ → CY mutation. **B.** Diagnostic single ion chromatograms of signal corresponding to *At*PEN2 tryptic peptide containing the V^211^ residue, obtained during analysis of leaf samples from the indicated lines. **B.** Diagnostic single ion chromatograms of signal corresponding to PEN2_DSB tryptic dipeptide containing the DSB formed between the conserved C^202^ residue and the mutation-introduced Cys residue, obtained from the same leaf samples as above.

**Figure S8. Lack of the disuffide bond (DSB) is conserved among putative *At*PEN2, *At*BGLU27, and *Br*BABG orhologs and close homologs (BGLU27-like) within Brassicaceae family.** Multiple sequence alignment of regions corresponding to the two conserved Cys residues involved in DSB formation shown for orthologs and close homologs of the investigated BGLUs identified in selected Brassicaceae genomes (Fig. 7; Table S2).

**Figure S9. Lack of the disuffide bond (DSB) is conserved among putative *At*PEN2 othologs and *Br*BABG close homologs (BABG-like) identified in Capparaceae and Cleomaceae species.** Multiple sequence alignment of regions corresponding to the two conserved Cys residues involved in DSB formation shown for orthologs and close homologs of the investigated BGLUs identified in selected Capparaceae and Cleomaceae genomes (Fig. 7; Table S2).

