## Supporting Information for "Loss of a conserved disulfide bond defines Penetration2-related immune myrosinases in Brassicaceae"

**Table S1** List of oligonucleotides used in the study. All sequences presented in the 5'-3' orientation. Colours highlight restriction sites: red – *Asi*SI; blue – *Sbf*I, or their post-digestion fragments: green – *Hind*III; magenta – *Aat*I. *Italics* indicate 14 bp of the 5′ *AtPEN2-TA* CDS located upstream of the *Aat*II site. Underline marks mutation introduced in the *PEN2_DSB* CDS.

| **Oligonucleotides to form synthetic DNA insert with *Asi*SI and *Sbf*I restriction sites.** | | |
|---|---|---|
| Upper strain | pAGCTTGCGATCGCACAAGAAGACCTGCAGGC*GAAGACGATTCTTC* GACGT | |
| Bottom strain | pC*GAAGAATCGTCTTC*GCCTGCAGGTCTTCTTGTGCGATCGCA | |
| **Primers** | | |
| **Gene ID** | **Primer name** | **Sequence** |
| **Cloning of *AtBGLU18* and *AtBGLU27*** | | |
| *BGLU18* (*AT1G52400*) | BGLU18_F | GCTTGCGATCGCATGGCACATCTTCAAAGAACATTT CCTACTGAGATGTCAAAAGGTAGATTAAACTTCCCT GAAGGC |
| | BGLU18_R | TTCGCCTGCAGGTGGAAACTGTGGTTTGAGG |
| *BGLU27* (*AT3G60120*) | BGLU27_F | GCTTGCGATCGCGATGAAGAAAAACTCGTTCGGCC GATC |
| | BGLU27_R | CTCGCCTGCAGGTTCTTCTTCTCTCTTCAAG |
| **Site-directed mutagenesis to create *PEN2_DSB*** | | |
| *BGLU26/PEN2* (*AT2G44490*) | DSB_F | TGGCGCTAGT<u>TGCTAC</u>GCTGGAATGTCGG |
| | DSB _R | TTAACATACTTGGAGCAC |
| **qPCR analysis** | | |
| *Actin1* (*AT2G37620*) | Actin_F | CCGGTATTGTGCTGGATTCT |
| | Actin_R | AATTTCCCGCTCTGCTGTTG |
| *BGLU26/PEN2* (*AT2G44490*) | PEN2-TA_F | GAAGACGATTCTTCGACGTCTAAGA |
| | PEN2-TA_R | TCAATTATTAGCTCCTTTGAAGAACAGAG |

**Table S2** Number of putative PEN2, BABG and BGLU27 orthologs and close homologs (BABG-like and BGLU27-like) in the members of Brassicaceae, Cleomaceae and Capparaceae families. Column colours according with Fig. 7. Phylogenetic relations within Brassicaceae family according to Brassibase (Kiefer *et al.*, 2014; German *et al.*, 2023).

| Family | Subfamily | Supertribe | Tribe | Species | Abbreviation | PEN2 | BABG | BABG-like | BGLU27 | BGLU27-like |
|---|---|---|---|---|---|---|---|---|---|---|
| Brassicaceae | Aethionemoideae | | Aethionemeae | *Aethionema arabicum* | Aa | 1 | 1 | 0 | 1 | 0 |
| | Brassicoideae | Arabodae | Alysseae | *Alyssum linifolium* | Alyli | 1 | 1 | 0 | 2 | 1 |
| | | | Arabideae | *Arabis alpina* | AA | 1 | 0 | 0 | 1 | 0 |
| | | Brassicodae | Brassiceae | *Brassica oleracea* | Bol | 1 | 0 | 0 | 1 | 1 |
| | | | | *Brassica rapa* | Brara | 1 | 2 | 0 | 1 | 1 |
| | | | | *Cakile maritima* | Camar | 1 | 1 | 0 | 1 | 1 |
| | | | | *Crambe hispanica* | Crahi | 1 | 1 | 0 | 2 | 1 |
| | | | | *Eruca vesicaria* | Eruve | 2 | 2 | 0 | 3 | 2 |
| | | | | *Sinapis alba* | Sialb | 2 | 1 | 0 | 2 | 1 |
| | | | Eutremeae | *Eutrema salsugineum* | Thhalv | 1 | 1 | 0 | 1 | 1 |
| | | | Isatideae | *Isatis tinctoria* | Isati | 3 | 1 | 0 | 1 | 2 |
| | | | | *Myagrum perfoliatum* | Myper | 1 | 1 | 0 | 1 | 1 |
| | | | Thelypodieae | *Caulanthus amplexicaulis* | Caamp | 1 | 1 | 0 | 2 | 1 |
| | | | Thlaspideae | *Thlaspi arvense* | Thlar | 1 | 1 | 0 | 1 | 1 |
| | | | Schrenkielleae | *Schrenkiella parvula* | Sp | 1 | 1 | 0 | 1 | 1 |
| | | Camelinodae | Arabidopsideae | *Arabidopsis halleri* | Ah | 1 | 0 | 0 | 1 | 2 |
| | | | | *Arabidopsis lyrata* | AL | 1 | 0 | 0 | 1 | 1 |
| | | | | *Arabidopsis thaliana* | At | 1 | 0 | 0 | 1 | 0 |
| | | | Boechereae | *Boechera stricta* | Bostr | 1 | 0 | 0 | 1 | 1 |
| | | | Camelineae I | *Camelina sativa* | Csa | 0 | 0 | 0 | 2 | 0 |
| | | | | *Capsella grandiflora* | Cagra | 0 | 0 | 0 | 1 | 0 |
| | | | | *Capsella rubella* | Carub | 0 | 0 | 0 | 1 | 0 |
| | | | Cardamineae | *Cardamine hirsuta* | CARHR | 1 | 2 | 0 | 1 | 0 |
| | | | | *Rorippa islandica* | Roisl | 1 | 1 | 0 | 1 | 1 |
| | | | Descurainieae | *Descurainia sophioides* | Desop | 1 | 1 | 0 | 1 | 1 |
| | | | Lepidieae | *Lepidium sativum* | Lesat | 2 | 2 | 0 | 2 | 1 |
| | | | Malcolmieae | *Malcolmia maritima* | Mamar | 1 | 1 | 0 | 1 | 1 |
| | | Heliophilodae | Biscutelleae | *Lunaria annua* | Luann | 1 | 0 | 0 | 2 | 2 |
| | | | Iberideae | *Iberis amara* | Ibeam | 4 | 2 | 0 | 1 | 1 |
| | | Hesperodae | Chorisporeae | *Diptychocarpus strictus* | Distr | 1 | 0 | 0 | 1 | 0 |
| | | | Euclidieae | *Euclidium syriacum* | Eusyr | 1 | 1 | 0 | 1 | 0 |
| Capparaceae | | | | *Capparis spinosa* | Cs | 2 | 0 | 1 | 0 | 0 |
| Cleomaceae | | | | *Cleome violacea* | Clevi | 1 | 0 | 2 | 0 | 0 |
| | | | | *Gynandropsis gynandra* | GG | 1 | 0 | 1 | 0 | 0 |
| | | | | *Tarenaya hassleriana* | Th | 1 | 0 | 4 | 0 | 0 |

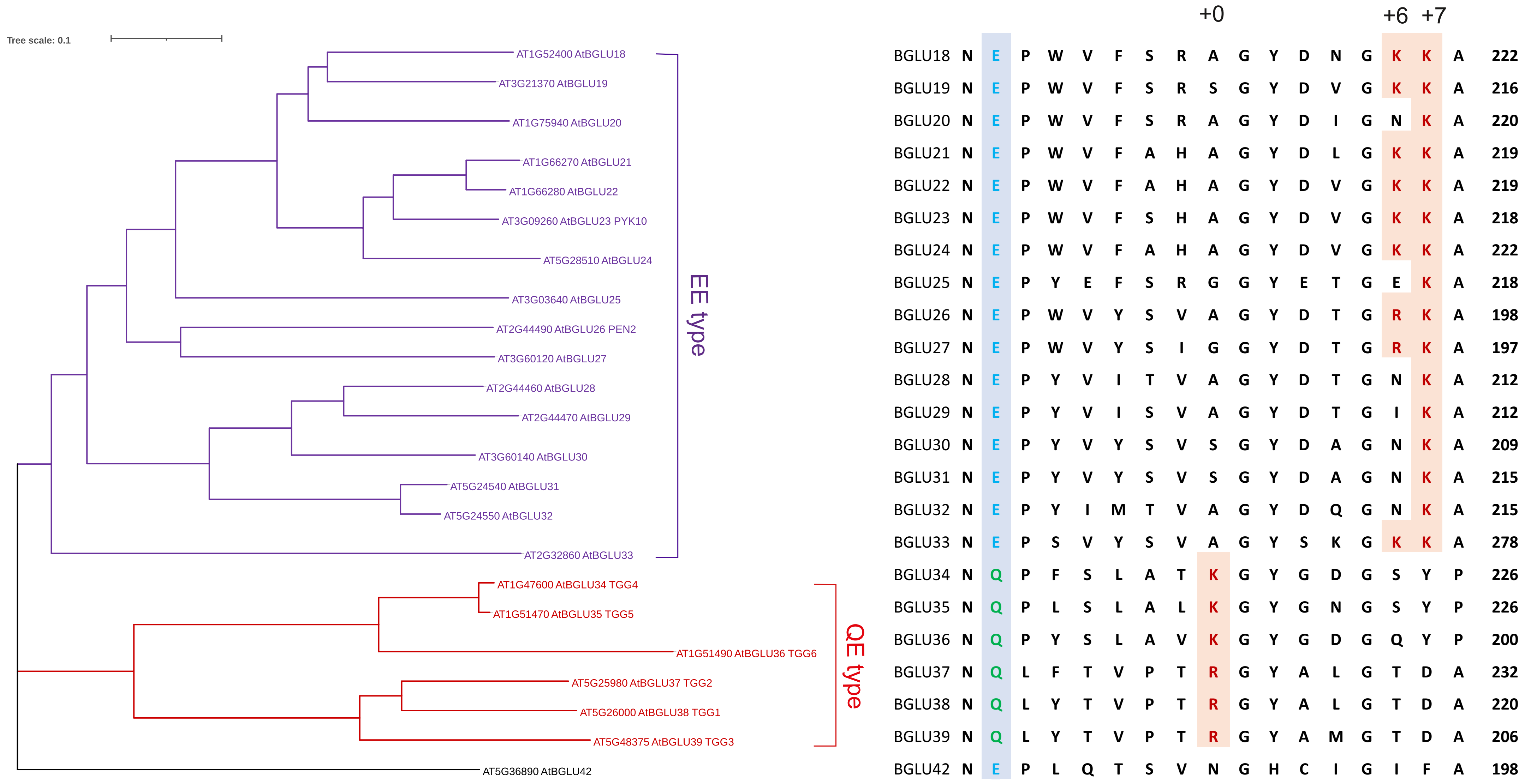

**Figure S1 QE- and EE-myrosinases in *Arabidopsis thaliana*.** The maximum-likelihood phylogenetic tree of AtBGLUs proposed to function as myrosinases. The positions of highlighted conserved catalytic (E/Q) and substrate recognition (RK) residues are based on Nakano *et al.* (2017).

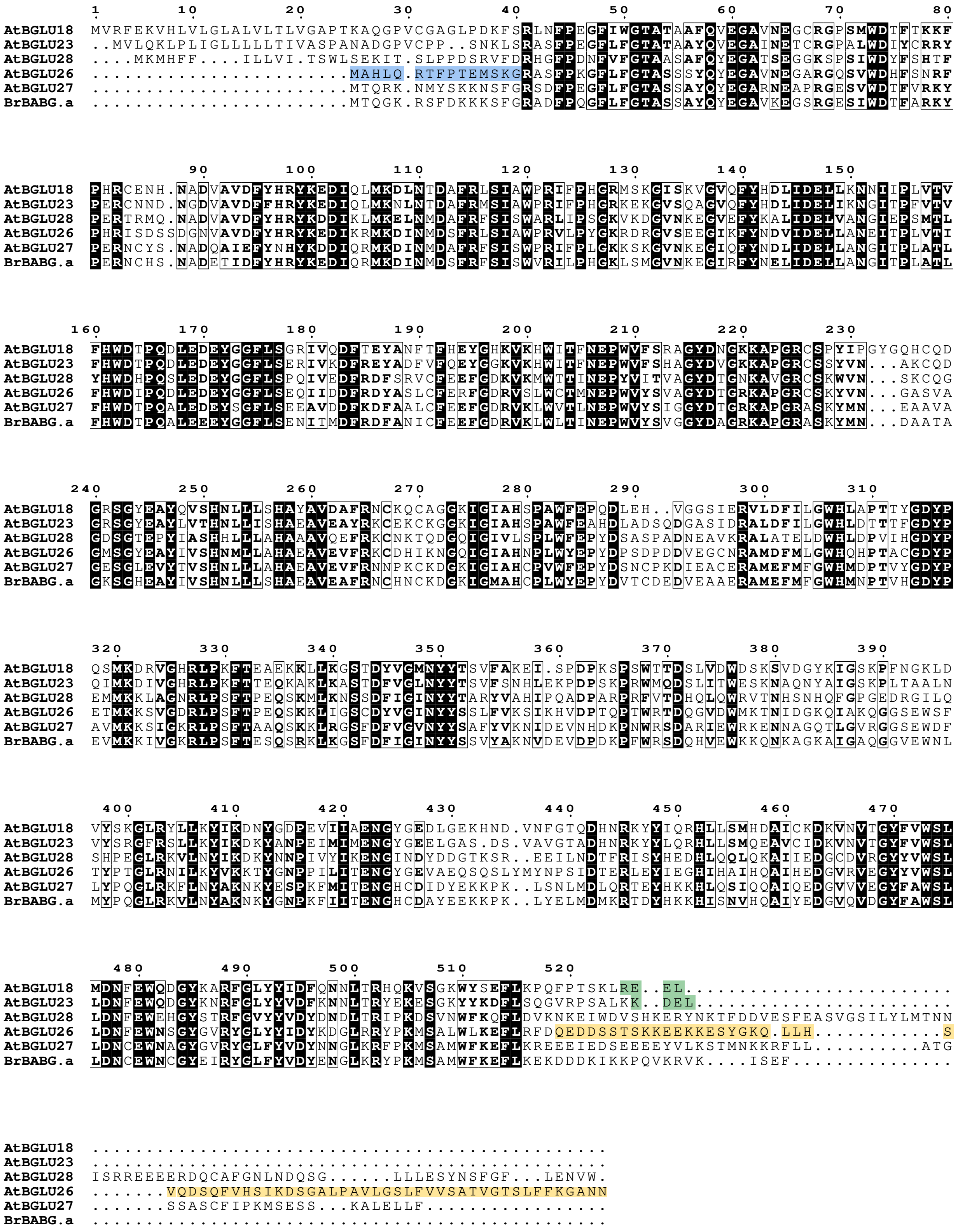

**Figure S2 Conservation of BGLUs selected for this study.** Protein sequence alignment of *At*BGLU26/PEN2 (AT2G44490), *At*BGLU18 (AT1G52400), *At*BGLU27 (AT3G60120), *Br*BABG.a (Brara.D02695), *At*BGLU23/PYK10 (AT3G09260) and *At*BGLU28 (AT2G44460) displaying conserved and variable regions among these myrosinases. The unique 15 aa N-terminal and 65 aa C-terminal regions of *At*PEN2 are underlined with blue and yellow colours, respectively. The C-terminal regions confining ER retention signal in *At*BGLU18 and *At*BGLU23 are highlighted in green colour.

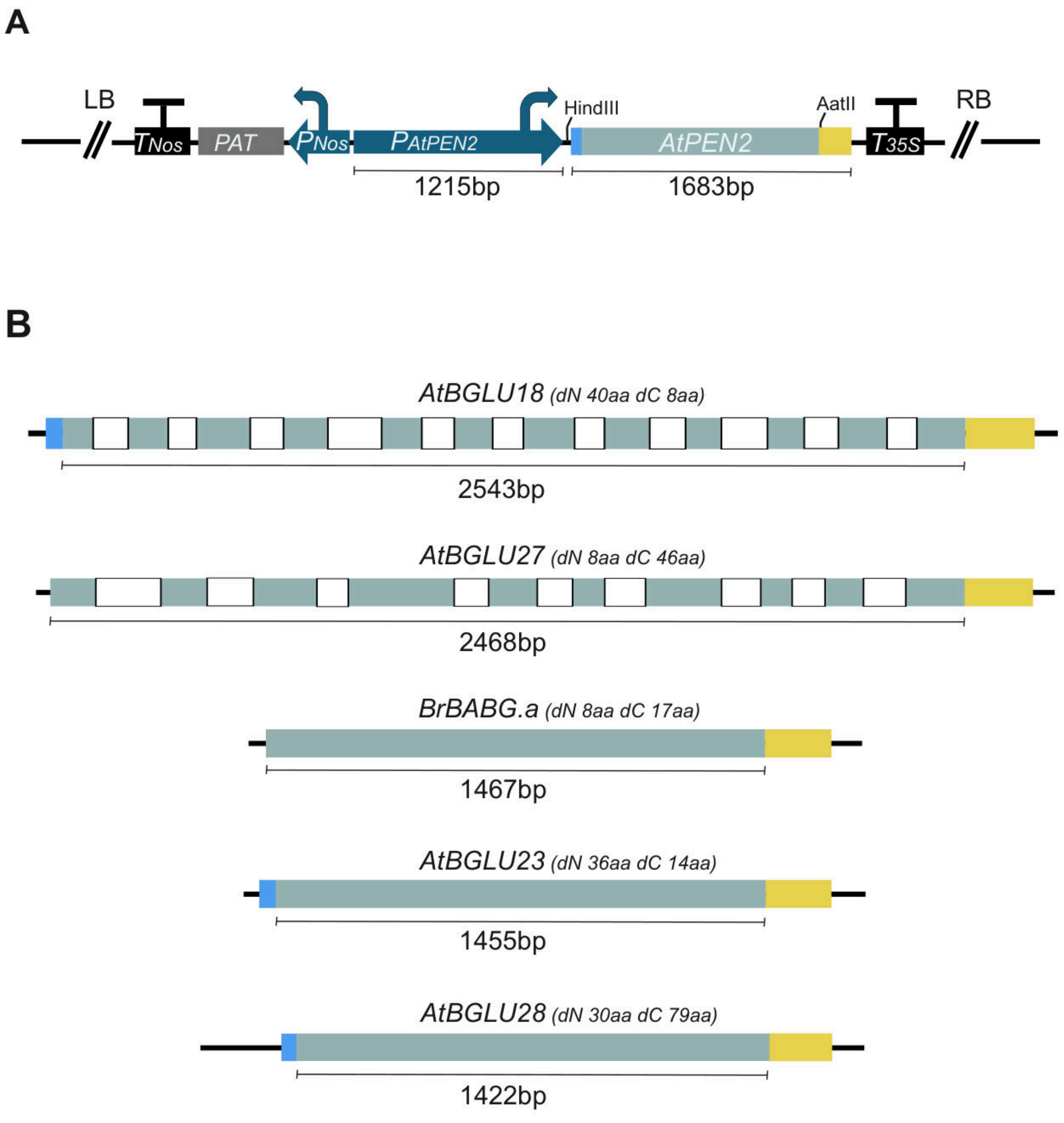

**Figure S3. Design of constructs used in this study. A**. Simplified exemplary map of the T-DNA fragment of the vector used for expression of *At*PEN2 under the control of its native promoter and for the preparation of the remining plasmids. **B**. Chimeric genes/CDS designed for expression of *AtBGLU26/PEN2* (AT2G44490), *AtBGLU18* (AT1G52400), *AtBGLU27* (AT3G60120), *BrBABG.a* (Brara.D02695), *AtBGLU23/PYK10* (AT3G09260) and *AtBGLU28* (AT2G44460).

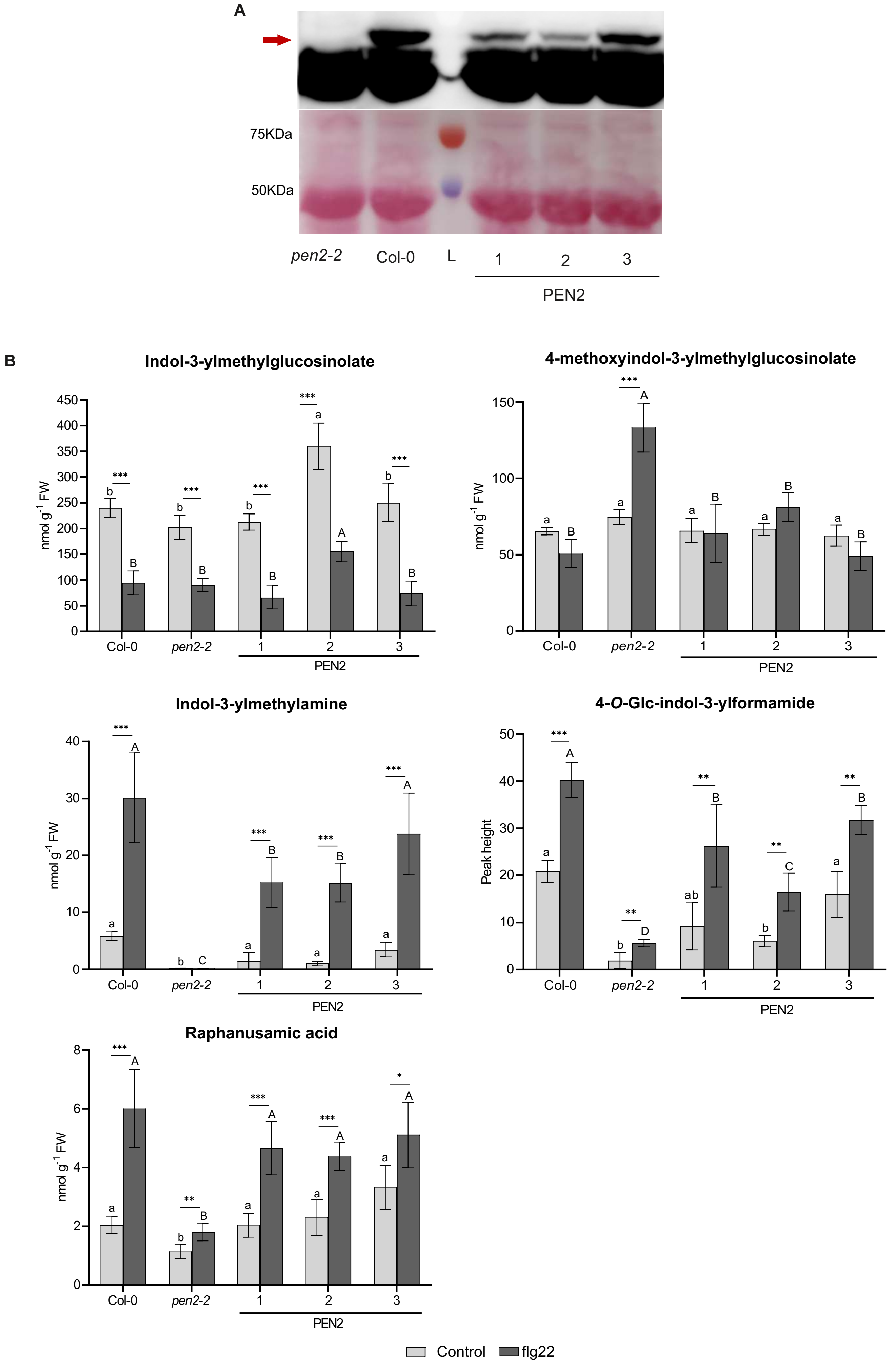

**Figure S4. Expression of *AtPEN2* in *pen2-2* mutant fully restores metabolic phenotype. A.** Immunoblot analysis of *At*PEN2 protein in leaf extracts from selected transgenic lines treated with flg22. Col-0 and *pen2-2* lines are included as positive and negative controls, respectively. The red arrow indicates position of *At*PEN2 (60 kDa) signal. **B.** Accumulation levels of selected indole glucosinolates and respective hydrolytic products in leaves of control and flg22-treated plants expressing *AtPEN2* construct. Presented results are means +/- SD. Significantly different statistical groups of genotypes indicated by the analyses of variance (ANOVA; $P < 0.05$; Tukey's post-hoc test) are shown with lower case (control) and upper case (flg22) letters. Values marked with asterisks are significantly different from respective controls (Student's unpaired t-test; * $P < 0.05$, ** $P < 0.01$, *** $P < 0.001$). FW - fresh weight.

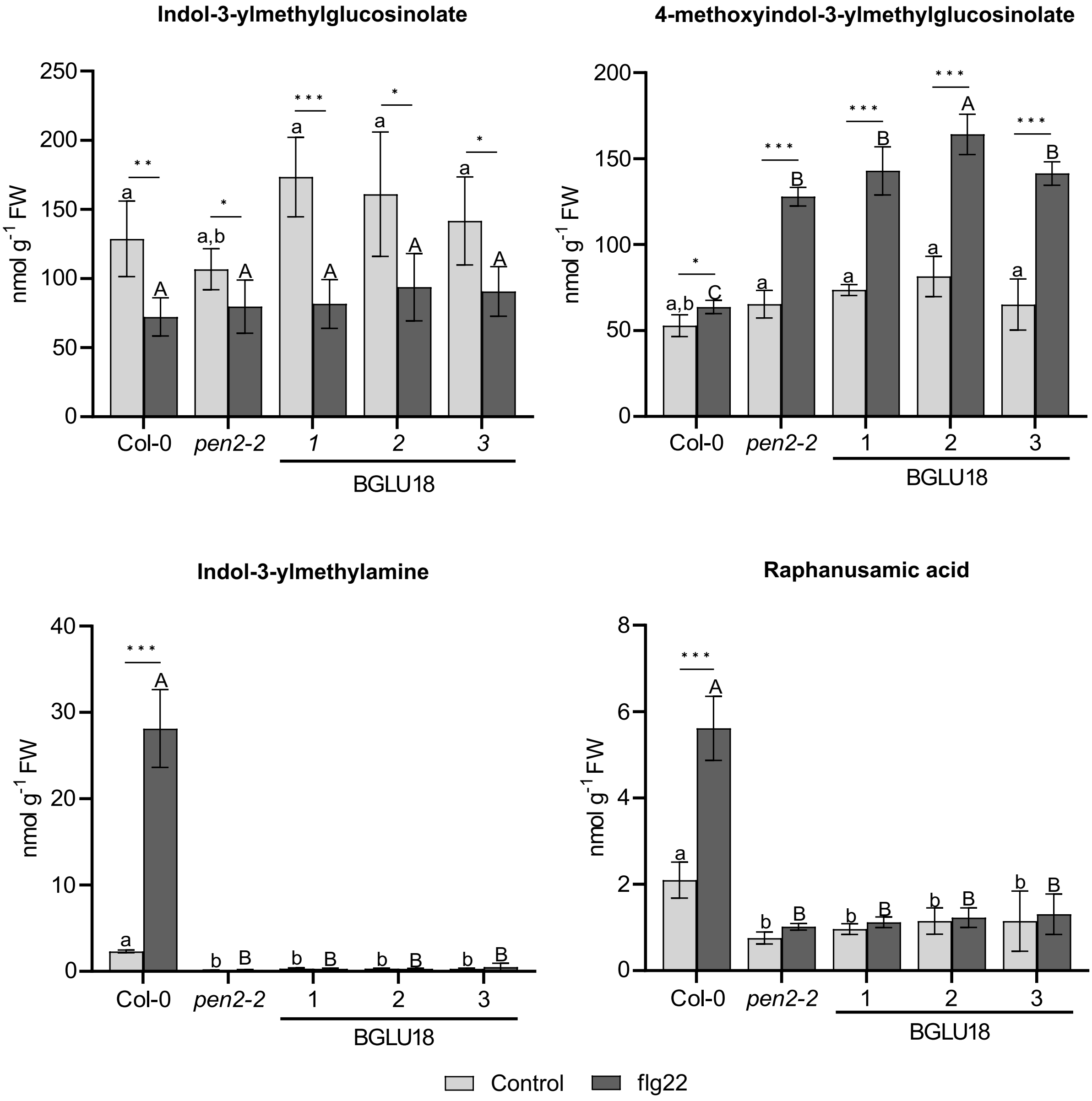

**Figure S5. Expression of *AtBGLU18* construct does not alter accumulation of indole glucosinolates and related hydrolytic products.** Accumulation levels of selected indole glucosinolates and respective hydrolytic products in leaves of control and flg22-treted plants expressing chimeric *AtBGLU18*. Col-0 and *pen2-2* lines are included as positive and negative controls, respectively. Presented results are means +/- SD. Significantly different statistical groups of genotypes indicated by the analyses of variance (ANOVA; $P < 0.05$; Tukey's post-hoc test) are shown with lower case (control) and upper case (flg22) letters. Values marked with asterisks are significantly different from respective controls (Student's unpaired t-test; * $P < 0.05$, ** $P < 0.01$, *** $P < 0.001$). FW - fresh weight.

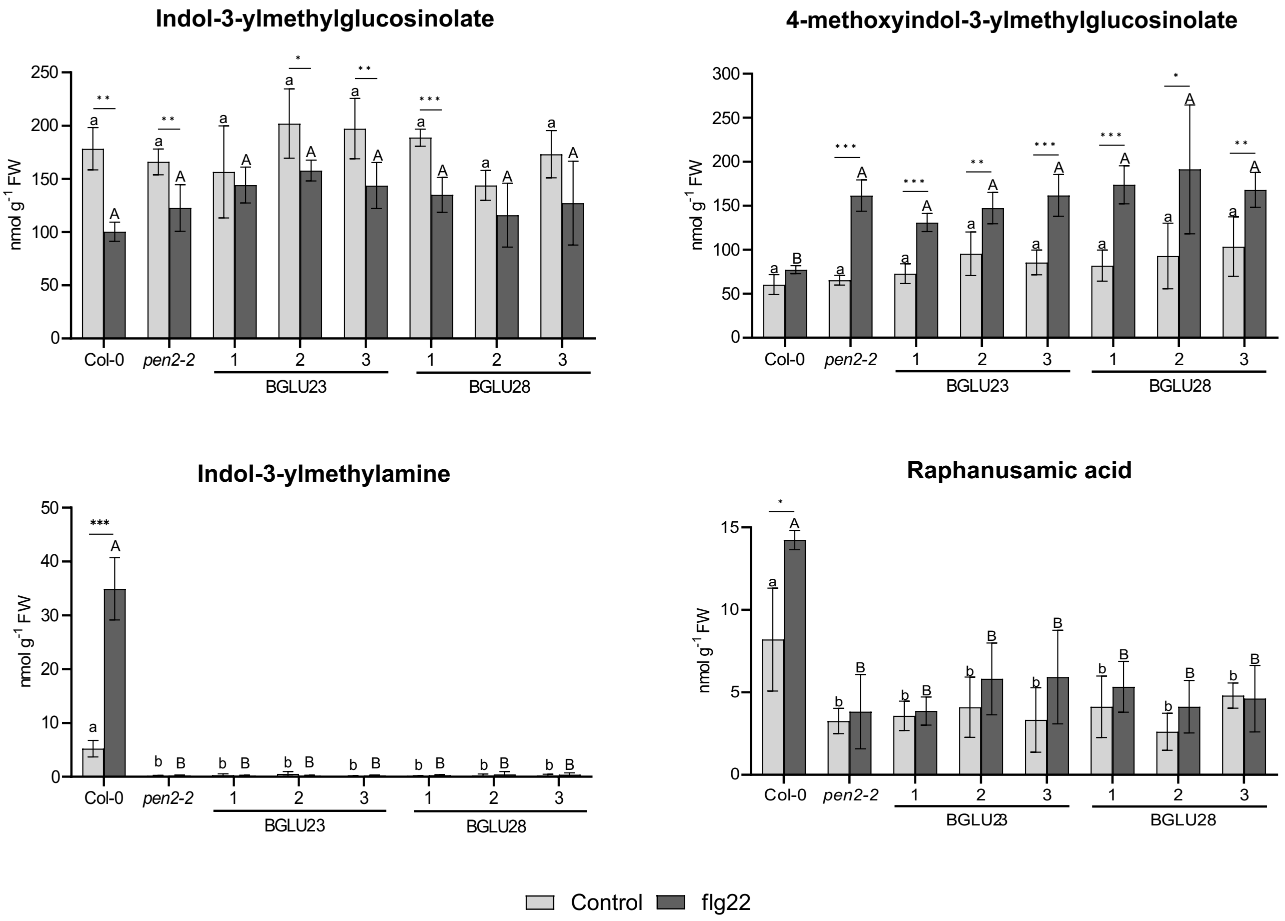

**Figure S6. Expression of *AtBGLU23* and *AtBGLU28* constructs does not alter accumulation of indole glucosinolates and related hydrolytic products.** Accumulation levels of selected indole glucosinolates and respective hydrolytic products in leaves of control and flg22-treted plants expressing chimeric *AtBGLU23* and *AtBGLU28*. Col-0 and *pen2-2* lines are included as positive and negative controls, respectively. Presented results are means +/- SD. Significantly different statistical groups of genotypes indicated by the analyses of variance (ANOVA; $P < 0.05$; Tukey's post-hoc test) are shown with lower case (control) and upper case (flg22) letters. Values marked with asterisks are significantly different from respective controls (Student's unpaired t-test; * $P < 0.05$, ** $P < 0.01$, *** $P < 0.001$). FW - fresh weight.

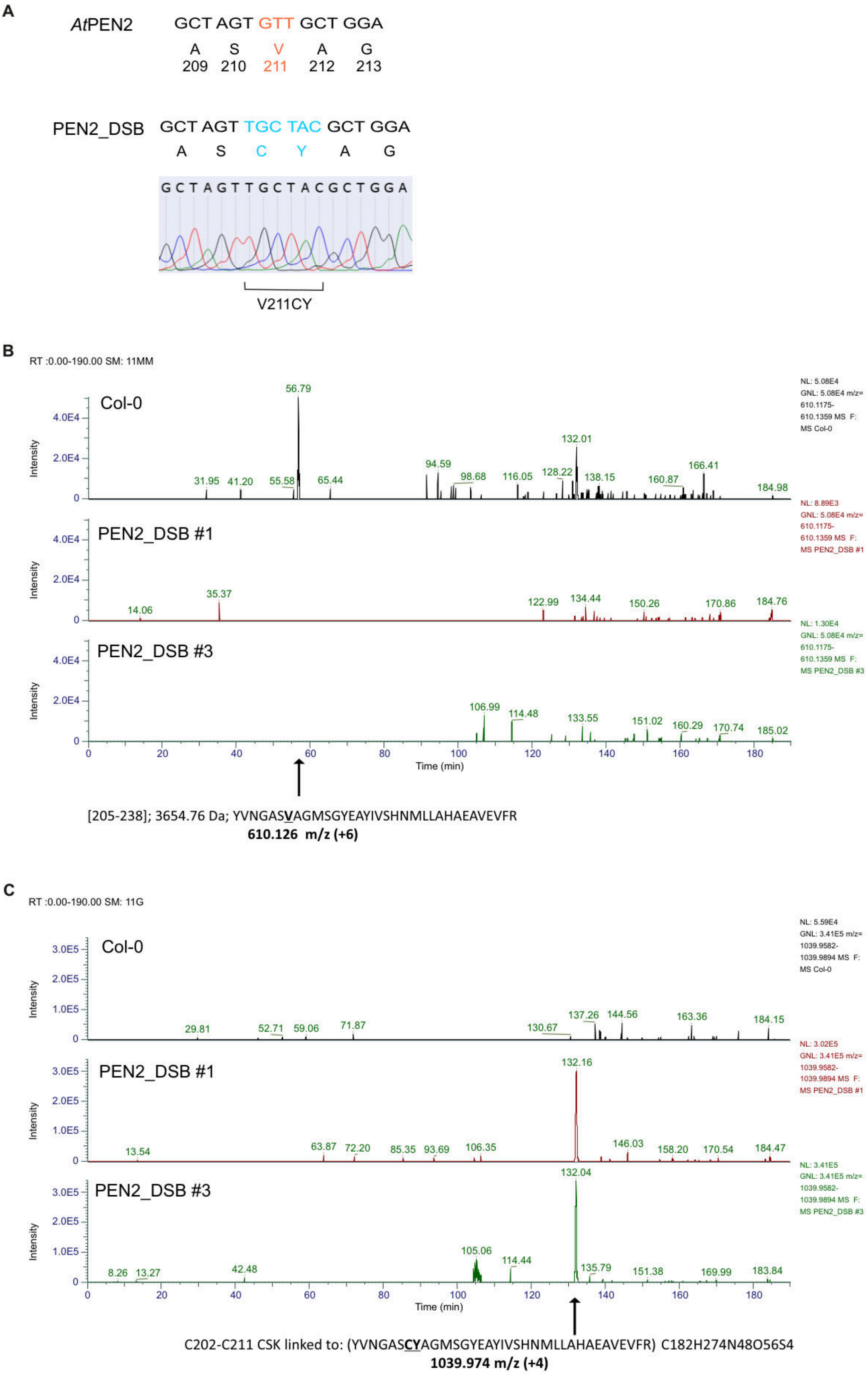

**Figure S7. Disulfide bond (DSB) is successfully restored in mutated *AtPEN2* protein (PEN2_DSB). A.** Sanger sequencing of the mutated *AtPEN2* CDS to validate $V^{211} \rightarrow$ CY mutation. **B.** Diagnostic single ion chromatograms of signal corresponding to *AtPEN2* tryptic peptide containing the $V^{211}$ residue, obtained during analysis of leaf samples from the indicated lines. **B.** Diagnostic single ion chromatograms of signal corresponding to PEN2_DSB tryptic dipeptide containing the DSB formed between the conserved $C^{202}$ residue and the mutation-introduced Cys residue, obtained from the same leaf samples as above.

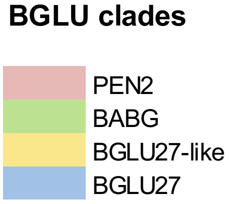

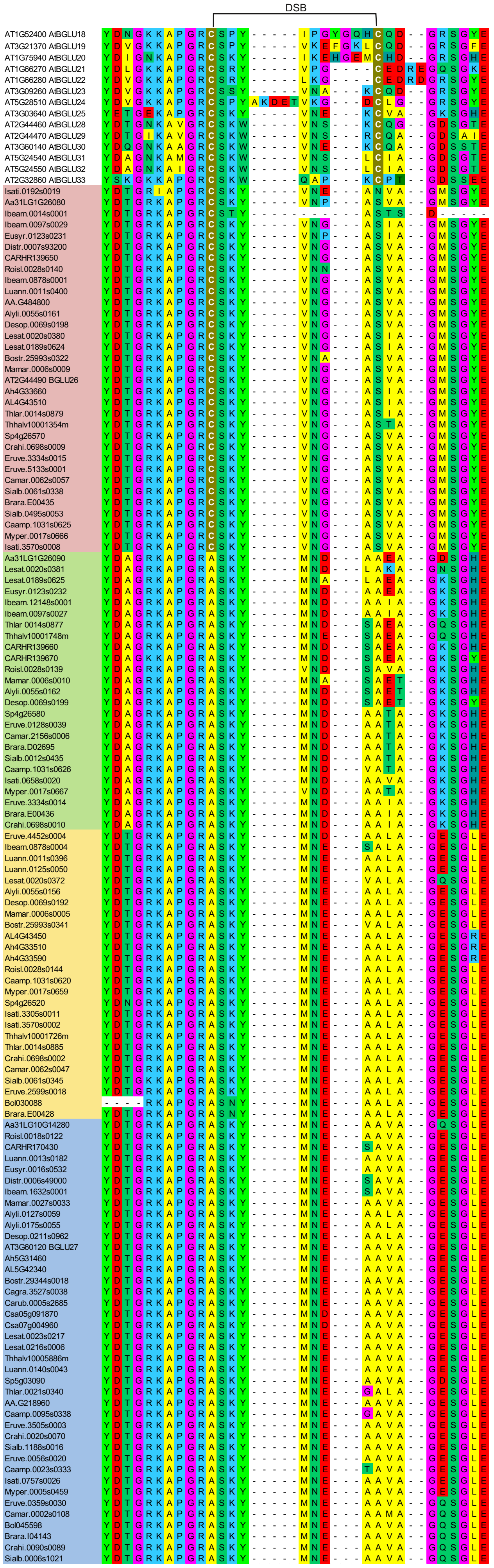

**Figure S8. Lack of the disulfide bond (DSB) is conserved among putative *At*PEN2, *At*BGLU27, and *Br*BABG orthologs and close homologs (BGLU27-like) within Brassicaceae family.** Multiple sequence alignment of regions corresponding to the two conserved Cys residues involved in DSB formation shown for orthologs and close homologs of the investigated BGLUs identified in selected Brassicaceae genomes (Fig. 7; Table S2).

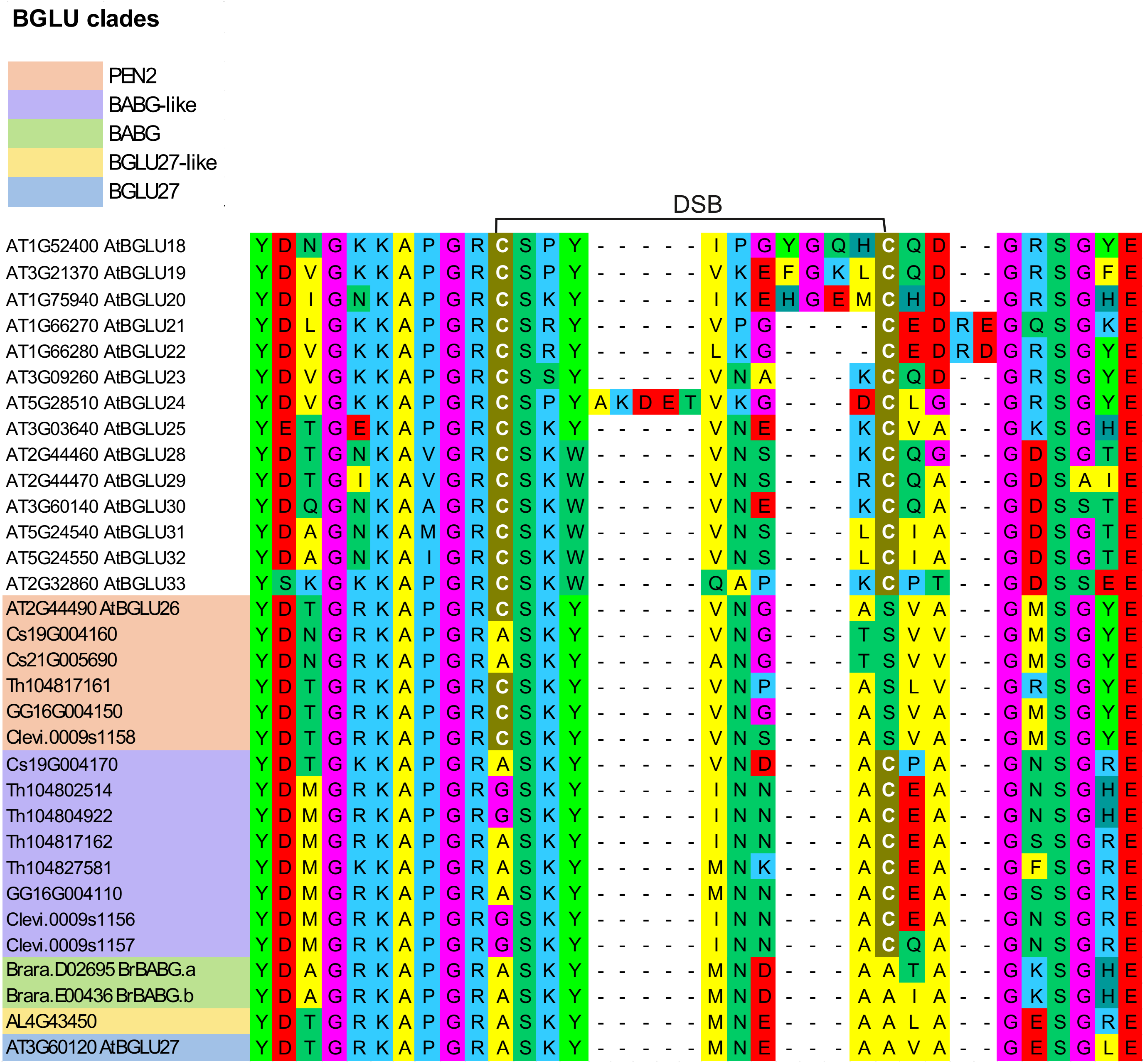

**Figure S9. Lack of the disuffide bond (DSB) is conserved among putative *AtPEN2* othologs and *BrBABG* close homologs (BABG-like) identified in Capparaceae and Cleomaceae species.** Multiple sequence alignment of regions corresponding to the two conserved Cys residues involved in DSB formation shown for orthologs and close homologs of the investigated BGLUs identified in selected Capparaceae and Cleomaceae genomes (Fig. 7; Table S2).
